# Long-range phase synchronization of high-gamma activity in human cortex

**DOI:** 10.1101/442251

**Authors:** G Arnulfo, SH Wang, B Toselli, N Williams, J Hirvonen, MM Fato, L Nobili, F Cardinale, A Rubino, A Zhigalov, S Palva, JM Palva

## Abstract

Inter-areal synchronization of neuronal oscillations below 100 Hz is ubiquitous in cortical circuitry and thought to regulate neuronal communication. In contrast, faster activities are generally considered to be exclusively local-circuit phenomena. We show with human intracerebral recordings that 100–300 Hz high-gamma activity (HGA) may be synchronized between widely distributed regions. HGA synchronization was not attributable to artefacts or to epileptic pathophysiology. Instead, HGA synchronization exhibited a reliable cortical connectivity and community structures, and a laminar profile opposite to that of lower frequencies. Importantly, HGA synchronization among functional brain systems during non-REM sleep was distinct from that in resting state. Moreover, HGA synchronization was transiently enhanced for correctly inhibited responses in a Go/NoGo task. These findings show that HGA synchronization constitutes a new, functionally significant form of neuronal spike-timing relationships in brain activity. We suggest that HGA synchronization reflects the temporal microstructure of spiking-based neuronal communication *per se* in cortical circuits.

## Introduction

Observations of organized neuronal population activities at frequencies above 100 Hz, such as high-gamma activity (HGA, 100–300 Hz)^1^, high-frequency oscillations (HFOs, 100–200 Hz) and ripple oscillations (150–200 Hz)^2, 3^ are abundant in electrophysiological recordings of both humans^1, 4^ and animals^5, 6^. HGA has been linked with perceptual and cognitive processes^1, 4, 7, 8^. Overall, high-amplitude HGA is a key signature of “active” neuronal processing. Ripple oscillations have been traditionally associated with sharp waves and off-task memory formation, but recent studies report ripples also during the performance of attention tasks^9^ and successful retrieval of memories^10^.

Electrophysiological HGA signals are thought to mainly arise from broad-band multi-unit spiking activity (MUA) and hence to directly reflect the local peri-electrode neuronal population activity *per se*^11, 12^. HGA^13^ and ripple oscillation signals do, however, also contain contributions from post-synaptic potentials. While the synaptic mechanisms underlying the hippocampal ripple oscillations are already well understood, it appears that genuine oscillations with presumably synaptic-communication-based mechanisms also contribute to the HGA signals^13^.

HGA has been thought to be exclusively local in terms of spike synchronization and phase coupling. For slower (< 100 Hz) neuronal oscillations, phase relationships among distributed neuronal assemblies are instrumental for coordinating neuronal communication and processing^14, 15^. Several lines of experimental and theoretical evidence have shown that these phase relations are dependent on frequency and neuroanatomical distance, and on the axonal conduction delays in particular, so that slow oscillations are generally more readily phase coupled over long distances than fast oscillations^16–18^. In line with this notion, measurements of long-range phase coupling in animal^15, 19^ and human^16^ brains suggest that neuronal oscillations only in frequency bands below 100 Hz exhibit inter-areal phase synchronization^20^ whereas the inter-areal cooperative mechanisms of HGA have remained poorly understood.

We hypothesized that long-range synchronization and phase coupling of HGA, even if unexpected, could nevertheless be conceivable because local synchronization and high collective firing rate can endow local pyramidal cell populations greatly enhanced efficiency in engaging their post-synaptic targets^21^, which would be experimentally observable as inter-areal HGA phase coupling. Such a finding would constitute a direct indication of spiking-based long-range neuronal communication *per se*. Nevertheless, there is little evidence for HGA synchronization over long distances so far^10, 22^.

In this study, we searched for long-range HGA synchronization using an extensive database of resting-state human stereo-electroencephalography (SEEG) recordings (*N* = 67). We used sub-millimeter accurate SEEG-electrode localization^23^ and white-matter referencing^16^ to obtain neocortical local-field potential (LFP) signals with little distortion from signal mixing with neighboring grey-matter or distant volume-conducted sources. We found that among these LFPs, long-range HGA synchronization was a surprisingly robust phenomenon and much stronger than synchronization at around 100 Hz. We rigorously excluded the possibilities of HGA synchronization being attributable to putative confounders such as the epileptiform pathophysiology or physiological and technical artefacts. The network architecture of resting-state functional connectivity achieved by HGA synchronization was split-cohort reliable and had a modular large-scale architecture that was distinct from those of lower frequencies. Finally, as two lines of evidence for functional significance, we found HGA synchronization to exhibit distinct cortical structures during sleep and awake states as well as to reflect the large-scale cortical processing underlying response inhibition in a visual Go/NoGo task. These findings thus reveal in the human brains a novel functionally-significant form of spatio-temporally highly accurate neuronal coupling, which elucidates the cerebral organization of neuronal communication.

## Results

### Probing human large-scale brain dynamics with SEEG

We recorded ∼10 minute resting-state human cortical (local-field potential) LFP signals from 92 consecutive patients with stereo-electroencephalography (SEEG). Among them, 25 subjects were excluded from further analyses due to previous brain surgery (temporal lobotomy) or an MRI-identified cortical malformation (Suppl. Table 1). The final cohort of 67 patients yielded a total of 7068 non-epileptic grey matter contacts (113 ± 16.2 per subject, mean ± SD, range 70-152) that gave a dense sampling across all neocortical regions (Fig. 1a) and of seven canonical functional brain systems defined by fMRI intrinsic connectivity mapping^24, 25^ (Fig. 1b, Suppl. Fig. 1).

**Figure 1.**
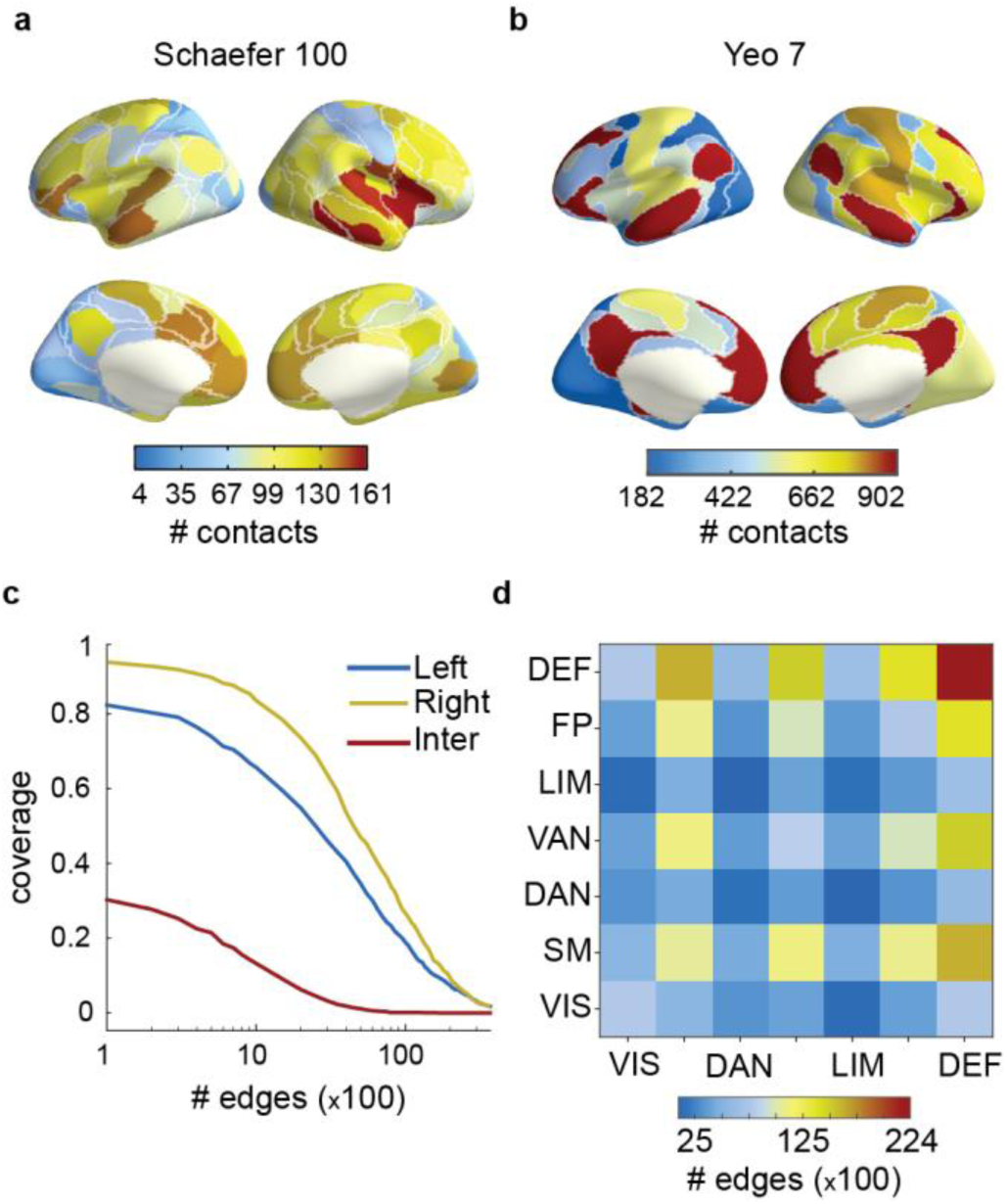
Anatomical sampling statistics and coverage of cortical interactions. **a,** Numbers of non-epileptic SEEG electrode contacts for each cortical area in a 100-parcel parcellation. **b,** Numbers of contacts in a parcellation of seven functional brain systems (see d). **c,** Cortical connectivity coverage in the 100-parcel parcellation for left-(blue), right-(yellow) and inter-hemispheric (red) region pairs connected by at least one electrode contact pair. **d,** Numbers of contact pairs connecting each pair of functional systems (visual, VIS; sensori-motor, SM; dorsal attention, DAN; ventral attention, VAN; limbic, LIM; fronto-parietal, FP; default mode, DEF).

We assessed the phase interactions between all LFP signals recorded from non-epileptic neocortical grey-matter locations. This yielded a dense coverage of cortical interactions with 5,500 ± 1,600 (mean ± SD) contact pairs per subject (range: 2,094–9,947) and a total of 368,043 contact pairs. Out of all possible within-hemispheric connections in the 100-parcel Schaefer atlas^25^, these recordings sampled at least 80% of the left- and 90% of the right-hemispheric connections (Fig. 1c) and provided abundant sampling on the scale of functional systems (Fig. 1d). The present data thus yield comprehensive insight into large-scale dynamics and connectivity in the human brain.

### Long-range phase correlations in high-gamma activity

We estimated inter-electrode-contact phase coupling with the phase-locking value (PLV) for 18 frequency bands between 3 and 320 Hz (see Methods). The inter-contact PLV estimates were averaged across subjects for three ranges of inter-contact-distance quartiles for each frequency band (Fig. 2a). The first quartile (0–2 cm) was excluded to avoid contamination from residual volume conduction. We found that the mean PLV increased from 3 to 7 Hz and then decayed rapidly from 10 to 100 Hz as expected^16^. However, from 100 Hz onward, the mean PLV increased monotonically, indicating highly significant HGA phase synchronization in all distance ranges.

**Figure 2.**
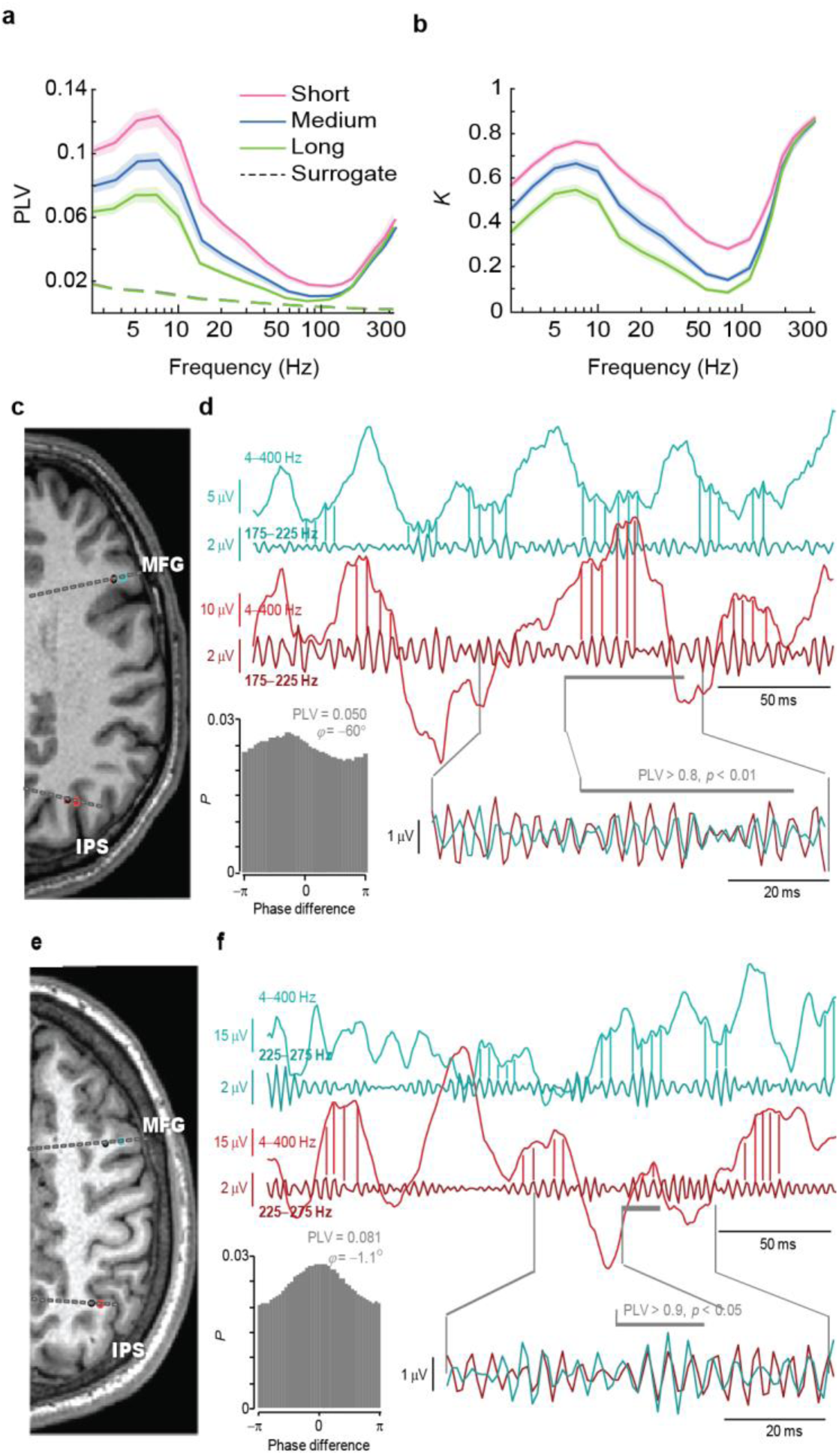
High-Gamma Activity shows robust long-range phase synchrony. **a,** Mean phase synchrony, measured with the phase-locking value (PLV) among all SEEG contacts and **b**, the fraction, *K*, of significant (*p* < 0.001) PLV observations for short (2–4.6 cm, pink), medium (4.6–6 cm, blue), and long (6–13 cm, green) distance ranges. Dashed lines represent the 99.9^th^ %-ile for surrogate data (*N_randomizations_* = 100). Shaded areas indicate the 2.5–97.5 %-ile bootstrap-confidence-limits for the mean PLV (*N_bootstraps_* = 100). **c-f,** Examples of long-range HGA phase synchrony with a non-zero phase lag (**c**, **d**) and a near-zero phase lag (**e**, **f**) in two subjects. Observation of the narrow-band HGA (175–225 Hz) peaks in the broad-band (4–400 Hz) time series shows that the narrow-band activity does not arise from spike or other filtering artefacts in these examples. MFG (azure traces), medial frontal gyrus; IPS (red traces), intra-parietal sulcus. In the MRI insets (**c, e**), the black dots represent the white-matter reference channel for each cortical grey-matter channel.

To quantify the neuro-anatomical extent of HGA synchronization, we assessed the connection density, *K*, that was defined as the fraction of contact pairs exhibiting significant (*p* < 0.001 for observed > surrogate PLV) phase synchronization (Fig. 2b). Even at long ranges (> 6 cm), more than 70% of >100 Hz connections were significant and both the PLV and *K* findings were split-cohort reliable (Suppl. Fig. 4). HGA phase synchronization was thus a widespread phenomenon in SEEG recordings of resting-state brain activity.

Visual inspection of HGA synchronization in SEEG electrode time series (Fig. 2 c–f) was in line with these observations and readily revealed episodes of significant HGA coupling in time windows across centimeter-scale distances. Notably, synchronized HGA was observed as low-amplitude oscillation-like activity that was visible in the time series also without filtering, which provides first evidence for that HGA synchronization was not attributable to spikes or technical artefacts.

Given the novelty and unexpected nature of these observations, we performed a series of control analyses to exclude the possibilities that HGA synchronization were due to: referencing schemes and volume conduction (Suppl. Fig. 1 & 2), non-neuronal signal sources (Suppl. Fig. 3), inadequate filter attenuation or settings (Suppl. Fig. 5), amplifier noise or pathological neuronal activity, such as inter-ictal spikes, or contamination from muscular signals (Suppl. Fig. 6). The results of these analyses converged on the conclusion that the seemingly anomalous HGA synchronization can only be explained by true long-range synchronization between local cortical neuronal assemblies (Suppl. Text). This conclusion was further consolidated by the findings, as detailed below, that HGA synchronization was predominant in specific functional systems, and had a community structure and laminar connectivity profile that were distinct from those of slower activities, which are inconceivable for technical or physiological artefacts.

### Neuroanatomical localization of high-gamma synchronization

To characterize the neuro-anatomical features of HGA synchronization networks, we investigated large-scale phase coupling of HGA in two spatial resolutions. First on the level of functional systems^24^, robust HGA phase synchrony was found within and between all systems, but the strongest and most widespread phase synchrony was found between and within the default-mode (DEF) and limbic (LIM) systems (Fig. 3a,b). This observation is in line with the hypothesis that HGA synchronization reflects healthy patterns of neuronal communication that is dominated by activity in these systems, DEF in particular^26^, during resting state. The observed systems-level connectivity pattern was split-cohort reliable (Suppl. Fig. 4), which rules out biases of individual subsets with distinct aetiologies.

**Figure 3.**
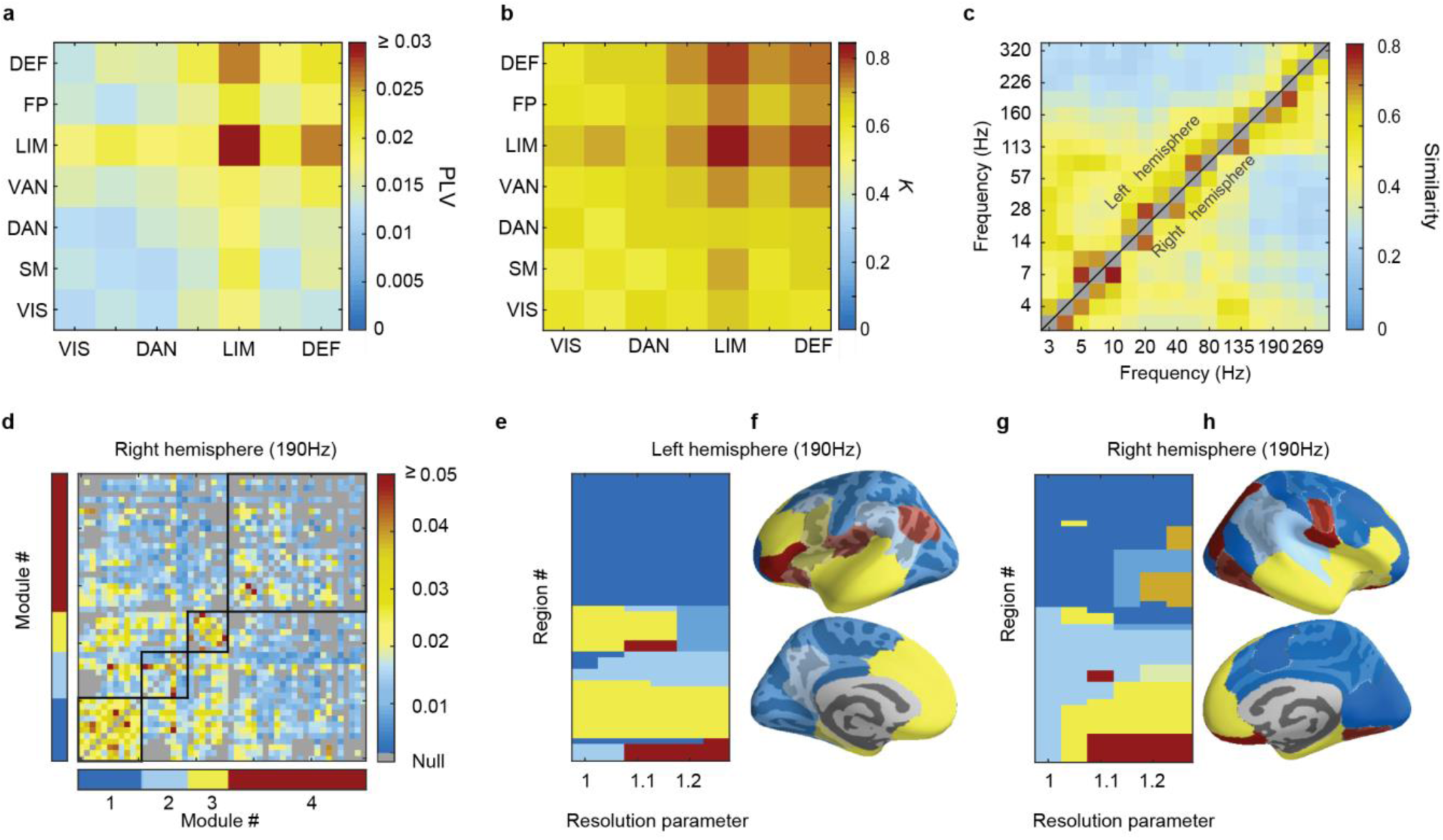
Neuroanatomical localization of high-gamma synchronization. **a,** Mean PLV across the HGA frequency band (113–320 Hz) among the seven functional brain systems. **b,** The fraction, *K*, of significant (*p* < 0.001) PLV observations in electrode contact pairs within (diagonal) or between (off-diagonal) the systems. **c,** Similarity of network module structures between different filter frequencies for 100-parcel resolution PLV connectomes, computed separately for the left and right hemispheres. Average similarity was computed with resolution parameter values from 1 to 1.25. **d,** With resolution parameter γ = 1.1, the Leiden method identified four modules in the 190 Hz PLV connectome for the right hemisphere (Null, unsampled connections). The allocation of left (**e**) and right hemisphere (**g**) cortical regions to modules in 190 Hz PLV matrix for resolution parameter values from 1 to 1.25. Neuroanatomical localization of cortical modules at γ = 1.1 for left (**f**) and right hemisphere (**h**) at 190 Hz (opaque color indicates significant, *p* < 0.05, parcel module-allocation stability).

To examine the architecture of HGA synchronization at a finer resolution, we used the 100-parcel Schaefer atlas^25^ and pooled data across subjects to estimate the connectome of inter-regional phase-synchrony for each frequency. We found these connectomes to be split-cohort reliable (permutation test, *p* < 0.05, one-tailed) across the range of frequency bands investigated, including the HGA frequency band (Suppl. Fig. 4). We then applied Leiden community detection to identify the putative communities therein and found sets of regions to represent robust modules in the HGA frequency bands (see Suppl. Text). The numbers of regions assigned reliably to communities were much higher than expected by chance (bootstrap test, *p* < 0.05, one-tailed) for each HGA frequency and throughout the investigated range of the Leiden resolution parameter, γ, values γ = 1–1.5 (Suppl. Fig. 9). These networks largely also demonstrated significant community structures compared to equivalent random networks (permutation test, *p* < 0.05, one-tailed) at a range of resolutions (Suppl. Fig. 9).

The communities in the high-gamma frequencies were similar to each other and dissimilar from communities in the lower frequency bands (Fig. 3c). For the 190 Hz connectome (Fig 3d), at a low spatial resolution, the network was split into four large communities that were divided into six to seven communities at higher resolutions (Fig. 3e, g). The majority of the communities comprised adjacent brain regions but also included spatially distant regions (Fig. 3f, h). Demarcation of spatially distal regions into the same modules is not only population level confirmation of long-range HGA synchrony observed at contact level, but also a strong indication of the functional relevance thereof. The whole-brain networks of HGA synchronization thus exhibited statistically significant and distinctive community structures in putatively healthy brain structures, as well as patterns of connection strengths that are stable across participants. These results thus further support the neurobiological origin of HGA synchronization.

### High-gamma phase correlations have a unique laminar profile

Deep and superficial cortical laminae are known to contribute differently to inter-areal phase-synchronization at frequencies below 100 Hz^16^. We next asked whether HGA synchronization networks also would show a similar differentiation between the laminae across distances. Leveraging our accurate localization of the electrode contacts, we divided the electrode contacts into “deep” and “superficial” by their cortical depth (see Methods), and assessed HGA phase coupling strength separately between the contacts in deep and superficial layers. This analysis reproduced the prior observation^16^ of low-frequency (< 30 Hz) phase coupling being stronger among superficial sources, which is well in line with recent observations about the laminar localization of, *e.g.*, alpha oscillation sources in human cortex^27^. In contrast, however, HGA synchronization was stronger (Fig. 4a, see inset) and more prevalent (Fig. 4b) among signals from deep cortical layers in all distance ranges (*p* < 0.05, Bonferroni corrected with *N*_freq._ = 18). This reverse pattern was also observed with iPLV and in bipolar recordings (Suppl. Fig. 2). Although the resolution of SEEG is insufficient for further investigating the underlying neuronal generators of these oscillations, *e.g.*, with a current source density analysis, our results indicate that HGA synchronization originates in neuronal circuitry distinct from that producing the slower LFP oscillations.

**Figure 4.**
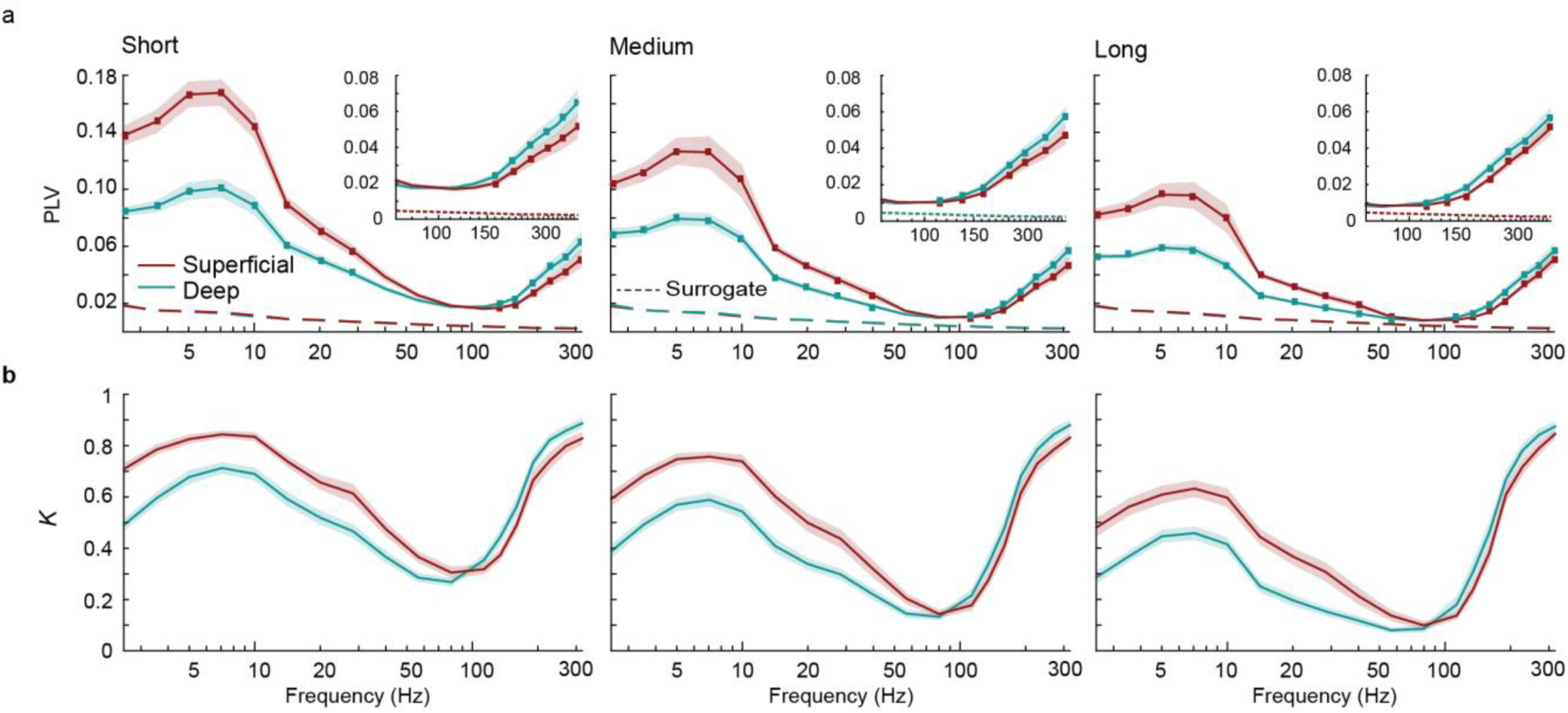
High-Gamma phase synchrony differs between cortical layers. **a,** Mean PLV at short (2–4.6 cm), medium (4.6–6 cm), and long (6–13 cm) distances exhibits distinct spectral profiles for SEEG electrode contact pairs located in deep cortical layers (−0.3 < GMPI ≤ 0; blue) and contact pairs in superficial cortical layers (0.5 < GMPI < 1; red). Dots represent the frequencies at which the difference between the superficial and deep contacts was significant (permutation test, *N_permutations_* = 100: *p* < 0.05, Bonferroni corrected for 18 frequencies). The GMPI value indicates the normalized depth of the electrode contact (0, white-grey surface; 1 pial surface). Inset: mean PLV of superficial and deep-layer contacts in the HGA frequency range. Dashed lines represent the 99.9^th^ %-ile for surrogate data (*N_randomizations_* = 100). Shaded areas indicate the 2.5–97.5 %-ile bootstrap-confidence-limits for the mean PLV (*N_bootstraps_* = 100). **b,** Fractions, *K*, of significant (*p* < 0.001) contact pair PLV observations.

### Long-range HGA synchronization is associated with local synchronization indexed by HGA amplitude

To further delve into the physiological plausibility of HGA synchronization, we asked whether it was related to the moment-to-moment variability in the amplitudes of local HGA. HGA amplitude is likely to tightly reflect the number of neurons in the local assembly and the degree of their local synchronization, both of which are central to the ability of a local assembly to engage its post-synaptic targets effectively. We thus hypothesize that the moments of strongest inter-areal HGA phase synchrony would coincide with the moments when both contacts recorded high HGA amplitudes. We selected electrode contacts exhibiting significant HGA synchronization and for each contact pair, distributed the data sample-by-sample into a 2D matrix according to quintiles of the normalized local amplitudes (see Methods). First, inspecting the variability in the numbers of samples among the cells of these 2D matrices we found that there was a slight positive correlation between the amplitudes so that the coincidence of the largest-quintile amplitudes was ∼1 % more prevalent than the coincidence of the smallest and largest amplitudes. Second, after equalizing the sample counts in amplitude quintile pairs, we estimated the PLV from samples in each quintile pair, and averaged the PLVs across electrode pairs and subjects for each frequency separately. We found that while HGA phase coupling was significant across a range of local amplitudes, it was the strongest in those moments when the HGA amplitudes were the largest in both contacts and essentially insignificant when either location exhibited the lowest HGA amplitudes (Fig. 5a, b). Quantitatively, PLV was very dependent on the local oscillation amplitudes and exhibited a ∼400 % difference between smallest and largest values in data averaged across the HGA band and a 200-800% difference in individual HGA frequencies with the strongest effects observed at highest frequencies (Suppl. Fig. 8). These data thus strongly support the notion of local HGA synchronization being instrumental for long-range HGA phase coupling.

**Figure 5.**
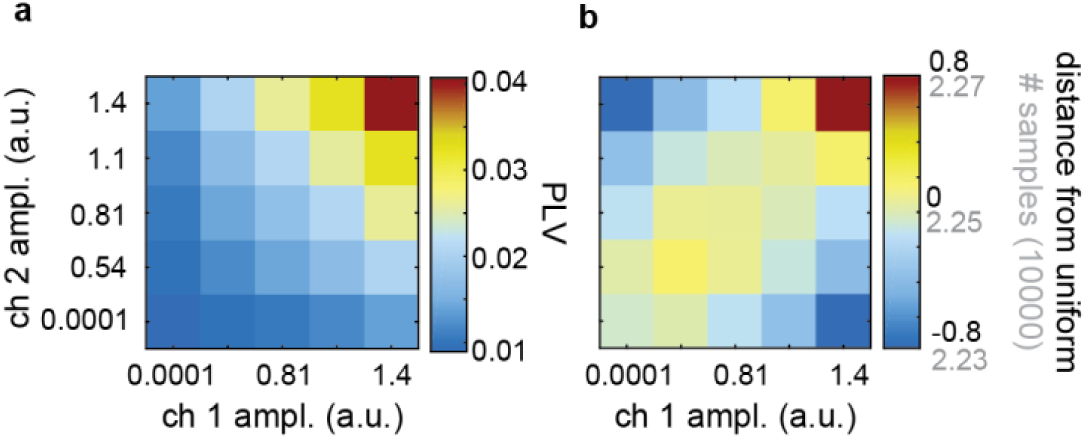
Stronger High-Gamma phase synchrony is correlated with maximal amplitude correlations between cortical sites. **a,** Moment-to-moment HGA synchronization (PLV) for SEEG electrode contact pairs (ch1, ch2) is dependent on the HGA amplitude at both contacts. Each matrix element is the mean of instantaneous PLV between all significant contact pairs (*p* < 0.001, *N_surrogates_* = 100) as a function of their moment-to-moment normalized amplitudes. **b,** HGA amplitudes are positively, albeit weakly, correlated. Numbers of samples (light grey) in each amplitude bin and their distance from a uniform distribution (black).

### Phase-amplitude coupling (PAC) of HGA with theta and alpha oscillations characterizes healthy brain dynamics

The “nesting” of fast oscillations in cycles of slower oscillations is an often-observed phenomenon in electrophysiological recordings at both < 1 Hz^28, 29^ and > 1Hz frequencies. We assessed whether HGA were nested within specific phase of slow rhythms by evaluating phase-amplitude correlations (see Methods) throughout the 3–320 Hz frequency range and among all pairs, excluding self-connections, of grey matter electrode contacts. We analyzed data from electrode contacts in the epileptogenic zone (EZ) to dissociate healthy from pathological patterns of PAC.

Among the putatively healthy LFP recordings, *i.e.*, nEZ contacts, the amplitudes of beta and low-gamma oscillations (14–40 Hz) were strongly coupled with the phase of theta and alpha oscillations (5–10 Hz) (Fig. 6a). Importantly, the amplitudes of HGA, peaking between 100–200 Hz, exhibited clear coupling with the phase of these theta–alpha oscillations. However, a fundamentally distinct cross-frequency PAC pattern was observed within the EZ (Fig. 6b), where instead of predominant narrow-band theta/alpha coupling, the amplitudes of oscillations from the lowest frequencies to HGA were locked to the phase of delta-band (here 3 Hz) oscillations. This putatively pathological pattern was also reflected in PAC evaluated between the healthy and epileptic areas, which exhibited both the delta and theta/alpha nesting of faster activities (Fig. 6c).

**Figure 6.**
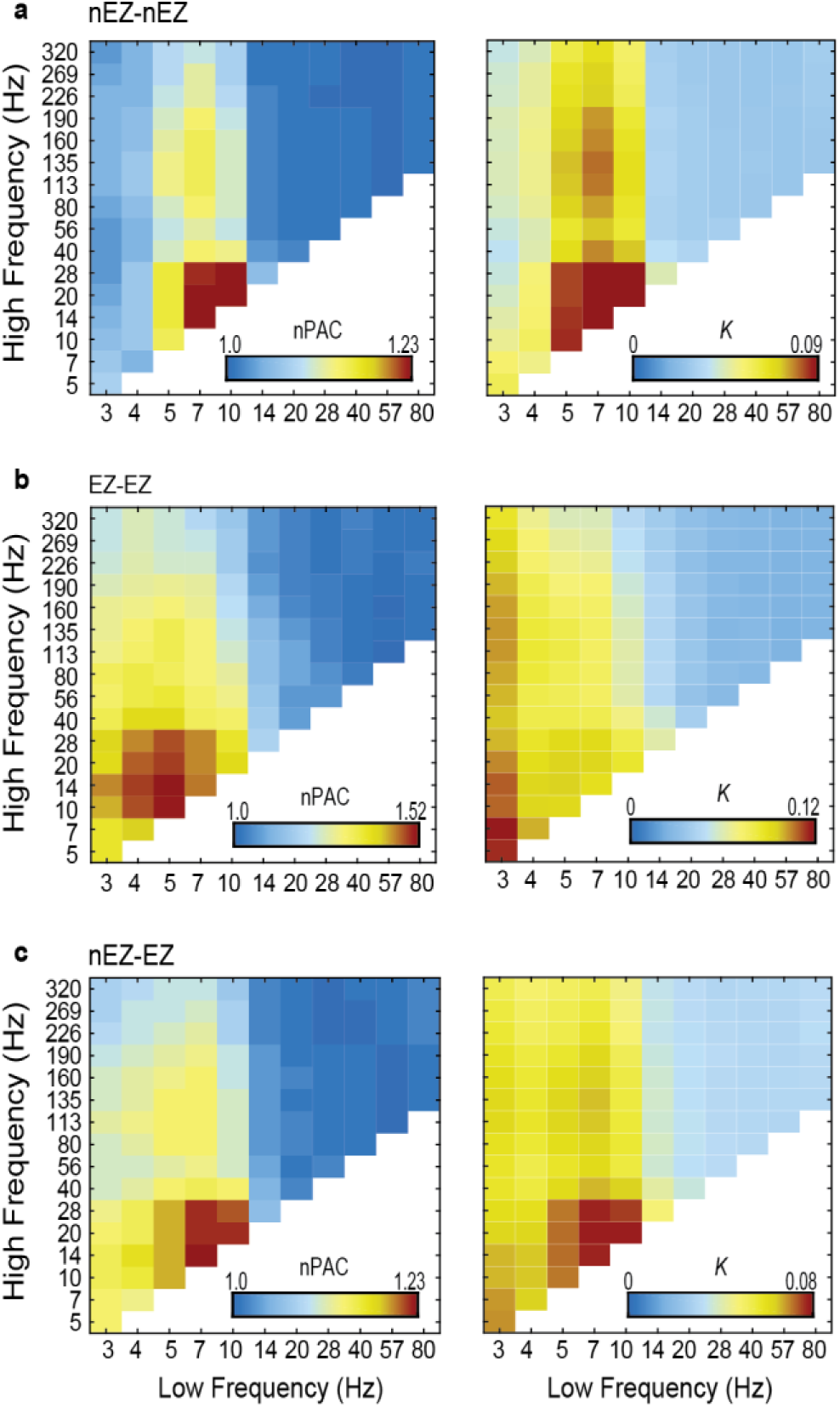
High-Gamma amplitude is modulated by the phase of slower oscillations. **a,** Among SEEG electrode contacts in putatively healthy brain regions (non-epileptogenic zone, nEZ), beta-to low-gamma band amplitudes (high frequency) are coupled with the phase of theta and alpha oscillations (low frequencies). nPAC: population mean of individual phase-amplitude-coupling (PAC) PLV estimates normalized by within-subject surrogate mean. Right: fraction of significant edges (*K*) for phase-amplitude coupling on the population level. **b,** Within the epileptogenic zone (EZ), prominent PAC is observed between the delta, theta, and alpha oscillations and the faster brain activities. **c,** PAC in connections between EZ and nEZ recording sites.

### Is high-gamma synchrony characteristic to the epileptic brain areas?

A crucial question for the functional implications of HGA synchronization is whether it is mainly a pathological property of the epileptogenic network or a feature of healthy brain activity and neuronal communication therein. The earlier finding of systematic HGA community structures across subjects with diverse aetiologies already provided evidence in support of the latter alternative. We further addressed this question first by asking whether the strength of HGA synchronization was significantly stronger in the EZ or between the EZ and nEZ. Controlling for the neuroanatomical distance of the electrode-electrode interactions, we found the epileptogenic zone to exhibit significantly stronger phase synchronization than healthy areas at low frequencies (3–10 Hz) but no significant differences were observed in the HGA frequency band (Fig. 7, see also Suppl. Fig. 6e).

**Figure 7.**
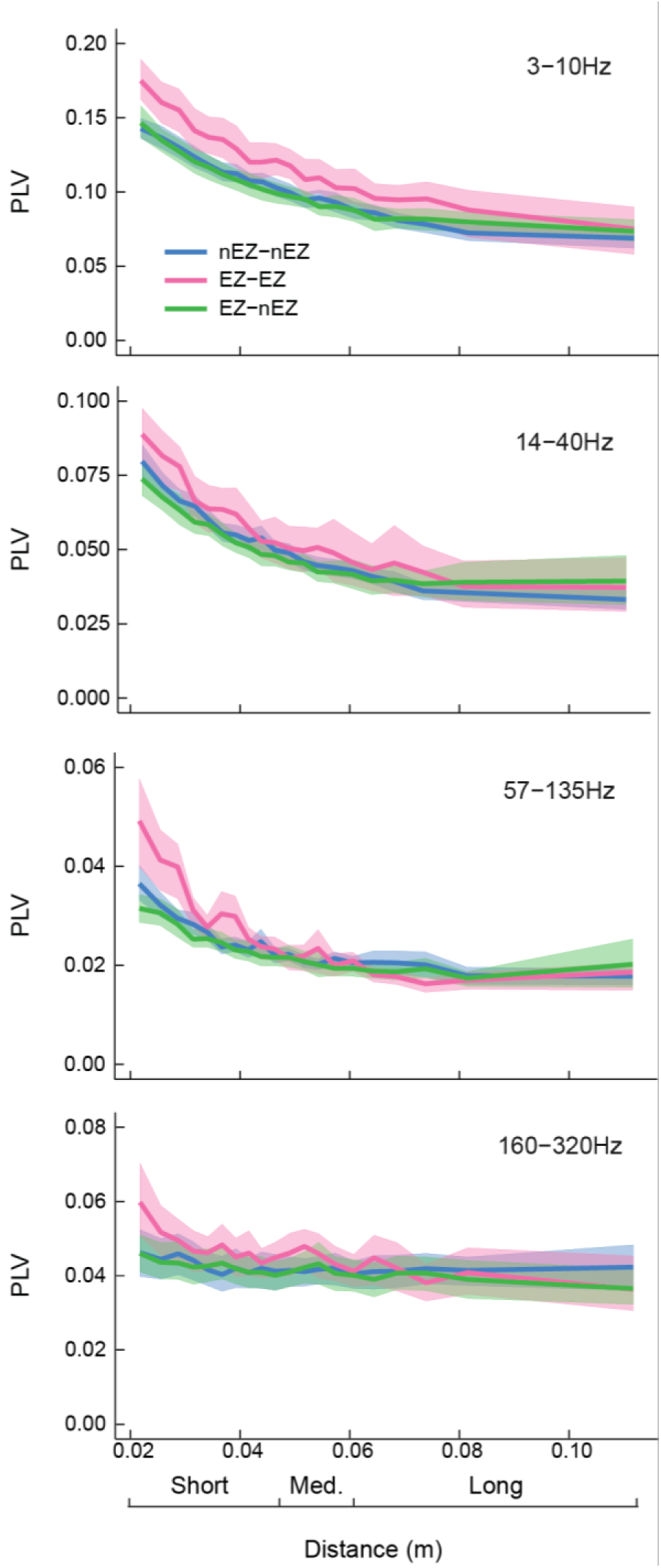
Phase synchronization is stronger within the epileptogenic zone (EZ) in low frequencies. At low frequencies (3–10Hz) and short-to-medium distances, mean phase synchrony among epileptogenic zone (EZ, pink) is significantly stronger than that among putatively healthy regions, *i.e.,* nEZ-nEZ (blue) or EZ-nEZ (green) connections. At higher frequencies, this difference between EZ-EZ and nEZ-nEZ and EZ-nEZ recording site is insignificant in all distance ranges. Shaded areas represent 5^th^ and 95^th^ percentile of PLV values (*N_bootstraps_* = 500).

We then tested whether the strength of HGA synchronization overall was correlated with the frequency of inter-ictal spikes; a proxy for the magnitude of epileptic pathology. No such correlation was observed (Suppl. Fig. 6). These data thus converged onto the conclusion that HGA synchronization is a property of healthy brain dynamics that is preserved in these patients.

### Distinct patterns HGA synchronization differentiate waking resting state and non-REM sleep

To address the putative physiological relevance of HGA synchronization, we first assessed whether it would reflect the large-scale brain state changes between awake resting state and sleep. We evaluated HGA phase synchrony with PLV in a subset of subjects (*n* = 7) with both resting-state and slow-wave sleep (SWS) recordings by pooling connections in the seven functional systems (see Fig. 3a, b). The wake and sleep system-level HGA synchronization matrices (Fig. 8a, b) were significantly different (*p* < 0.001, dissimilarity permutation test) so that during sleep, the pattern of HGA synchronization between systems was different, compared to during awake state. Further, increased HGA synchronization was found for sleep, within the limbic system and between the limbic and default-mode and ventral-attention systems (two-tailed t-test, *p <* 0.05, uncorrected). This finding as well as the anatomical pattern of system-system phase couplings was reproduced with iPLV (*p* < 0.001, dissimilarity permutation test, Supp. Fig. 7a, b). These findings are well in line with the notion of memory-consolidation-related 100–200 Hz ripple oscillations being a characteristic of non-REM sleep and indicate that such ripple activities may not only co-occur^10, 30^ but also exhibit systematic phase relationships within the limbic system and between the limbic and other functional systems.

**Figure 8.**
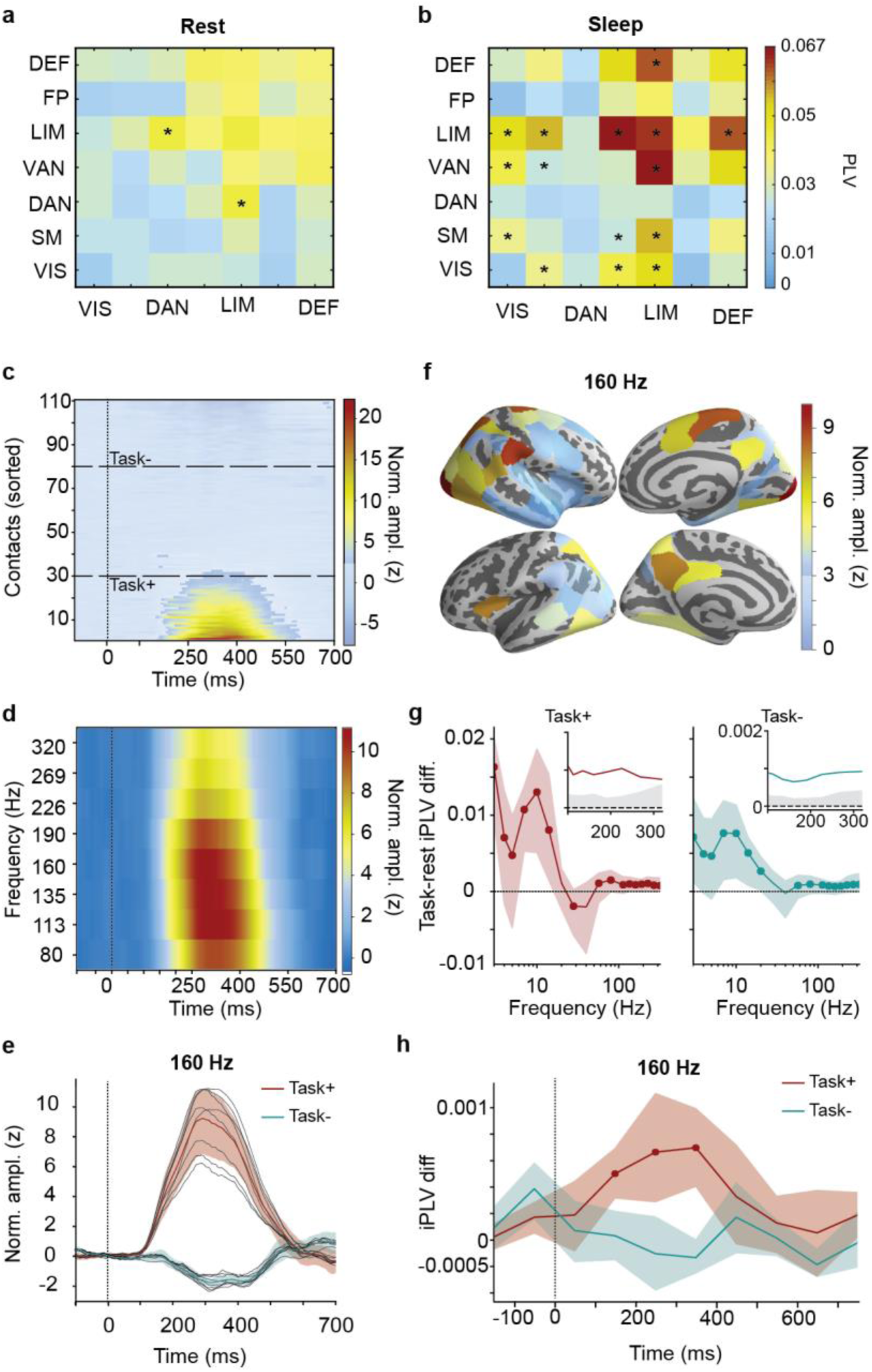
HGA synchronization is enhanced during sleep and visuomotor inhibition. PLV across high-gamma (HG) frequencies between systems are different for (**a**) rest and (**b**) sleep, (dissimilarity permutation test, *p* < 0.001, N=1,000, one-tailed conditions). In (**a**) asterisks indicate greater PLV during rest compared to sleep and in (**b**) asterisks indicate greater PLV during sleep (two-tailed post-hoc permutation test, *p* < 0.05, uncorrected). **c,** HGA amplitude increased above baseline after NoGo stimulus onset (at 0 ms) in a subset of contacts in a representative subject (averaged across 120–320 Hz, z-score normalized by the pre-stimulus baseline from −500 ms to −10 ms). Unshaded area indicates z > 3. **d,** HGA amplitude averaged across subjects and the top-30 task-positive contacts within subjects. **e,** Event-related response at 160 Hz averaged across subjects for the top-30 task-positive (red) and task-negative contacts (azure). Shaded areas indicate the 2.5–97.5 %-ile bootstrap-confidence-limits of the means (*N_bootstraps_* = 10000). **f,** Anatomical loci of the strongest event-related responses. **g,** Difference in HGA synchronization (iPLV) between task and rest. Colored shaded areas indicate confidence limits of the means (as in **e**), round markers indicate difference between task and rest (two-tailed permutation test). In insets, grey shades are 97.5 %-ile confidence limits for the null hypothesis of no difference (*p* < 0.05, two-tailed permutation test, *N_permutations_* = 10000). **h,** Time-resolved HGA synchrony (iPLV) among the top-30 task-positive (red) and task-negative contacts (azure) in response to the NoGo events. Shaded areas indicate the 2.5–97.5 %-ile bootstrap-confidence-limits of the means (*N_bootstraps_* = 10000). Round markers indicate time-windows where HGA synchronization in Task+ contacts is greater than among Task-contacts (*p* < 0.05, corrected, one-tailed max-statistic permutation test).

### Inter-areal HGA synchronization in the visual system is transiently enhanced during visuomotor processing

To elucidate the functional role of HGA synchronization beyond the insight yielded by its characteristics in resting-state activity and sleep-wake differences, we acquired resting- and task-state SEEG data from an additional cohort of 11 patients. The patients performed a visual Go/NoGo response-inhibition task where they reacted with a button press to Go stimuli (blue rectangles, 75% probability) and withheld responses to rare NoGo stimuli (yellow rectangles, 25% probability). In this task, correct response inhibition is known to involve large-scale fronto-parietal brain activation. We first examined the peri-stimulus amplitude dynamics of local HGA that has been established to indicate neuronal spiking in the immediate vicinity of the electrode contact and thereby to localize task-relevant brain areas with high accuracy^4^. In the latency range of 100–600 ms after the NoGo stimulus onset, we found 100–300 Hz HGA amplitude to increase above four baseline SDs (Fig. 8b) for at least 100 ms in 29–45 channels out of 103–139 grey matter recording sites in eight subjects. At the same latencies, markedly smaller HGA responses were observed for the Go stimuli (Suppl. Fig. 7c). Neither the latency nor duration of these responses were correlated with the reaction times suggesting that the HGA responses primarily reflect inhibition-related processing in this task. Three subjects showed no HGA responses and were excluded from further analyses. We used the 30 electrode contacts with the strongest amplitude effect for each subject to localize the cortical areas putatively underlying inhibition-relevant neuronal processing for the NoGo stimuli. Group-averaged time-frequency representation of these task-region amplitude data showed the amplitude effect was most pronounced in the 100-200 Hz frequency range (Fig. 8c), and at the peak frequency, individual HGA responses were clearly observable (136-184 Hz filter band, center frequency 160 Hz, Fig. 8d, see red and black lines). We also picked the 30 contacts with the smallest amplitude effect to localize the individual task-irrelevant or -negative brain areas and found them to exhibit a small amplitude decrease during this time window (Fig. 8d, blue line). The task-positive amplitude effects for the NoGo responses were predominantly localized into both ventral- and dorsal-stream visual areas and to the parietal cortex (Fig. 8e; see Suppl. Fig. 1 for sampling).

We first assessed the role of HGA synchronization in visual processing by comparing with time-averaged synchronization estimates of the Go/NoGo-task data with resting-state recordings acquired prior to the task. Among both the task-positive and -negative brain areas, HGA synchronization was stronger during task performance than during rest (Fig. 8e) and, importantly, HGA synchronization was significantly stronger among the task-positive than the -negative areas during task processing for all frequencies up to 160 Hz (*p* < 0.05, two-tailed permutation test, Benjamini-Hochberg FDR corrected). To address this in a time-resolved manner, we evaluated HGA synchronization in 100 ms time windows across trials. We also produced surrogate estimates with trial shuffling to exclude the putative contributions of stimulus-locked signal components and amplitude-change related changes in temporal correlation structures. We found HGA synchronization to be strengthened among the task-relevant brain areas during the period of active stimulus processing in response to NoGo stimuli so that the synchronization in the task-relevant areas was significantly stronger than in the task-irrelevant or -negative areas and significantly above the surrogates throughout the time windows from 50 to 450 ms (*p* < 0.05, max-value randomization test). The same analysis for the Go events revealed no significant increases in large-scale synchronization in any of the three groups. Moreover, HGA synchronization for the NoGo stimuli was significantly stronger than that for the Go stimuli in the 50-450 ms time range (*p* < 0.05, max-value randomization test, Suppl. Fig. 7d). These data thus show that HGA synchronization is strengthened from resting-state levels during task performance, associated with transient task-relevant neuronal processing, and specifically associated with the neuronal communication underlying the large-scale coordination of visuo-motor response inhibition in the Go/NoGo task.

## Discussion

HGA is thought to be a direct indication of ‘active’ neuronal processing *per se*^1, 4–6^ because of its strong correlation with neuronal firing rates and the BOLD signal^31, 32^, and with a range of cognitive processes^1, 4, 7, 8^. In the same frequency range, high-frequency oscillations (HFOs) are predominantly observed in epileptogenic brain areas and are elevated in magnitude prior to epileptic seizures^33^. The relationship between HGA and HGOs has remained unclear; HFOs may either be a phenomenon separate from HGA and integral to epileptic pathophysiology or, alternatively, pathological modulation of otherwise physiological HGA that also is known to exhibit oscillatory temporal patterning^13^.

In either case, HGAs or HFOs have been considered exclusively as local phenomena being phase-synchronized only within a local brain regions. HGA *amplitude* fluctuations may, however, be coupled inter-areally in human cortical networks during reading^34^, and similarly, ripple oscillations bursts co-occur between the rat hippocampus and association cortices during memory formation^30^. However, amplitude correlation does not have the temporal precision that phase-coupling has for carrying out important computational functions during neuronal processing. Nonetheless, inter-areal phase-coupling has been thought to exist only in slow brain rhythms but not in HGA frequency bands. To the best of our knowledge, there is only one reported HGA inter-areal *phase* coupling^22^. In this study, Yamamoto et al (2014) reported phase synchronization of ripple oscillations within the entorhinal-hippocampal circuit in behaving rats.

We report here that in the human brains, neuronal activity in the 100–300 Hz frequency range may be phase synchronized across several centimeters during awake resting-state brain activity. Long-range synchronization was highly reliable and not explained by physiological or technical artefacts in the recorded data. HGA synchronization was also primarily a physiological phenomenon and not explained by epileptic pathophysiology. Importantly, the large-scale patterns of HGA synchronization were distinct during awake resting state and non-REM sleep, indicating that HGA synchronization is dependent on global brain states. Specifically, this finding implies that the non-REM-sleep ripple oscillations entrain their post-synaptic targets in widespread brain areas, which may be essential for memory formation and consolidation^10^. Moreover, HGA synchronization was observed in a narrow time window specifically in response to stimuli for which the responses were correctly withheld, which strongly suggests that HGA synchronization is functionally significant in the neuronal communication underlying response inhibition in the present visual Go/NoGo task. HGA synchronization thus constitutes a novel and functionally relevant form of neuronal long-range coupling in the dynamic functional architecture on human brain dynamics. We suggest that these observations of HGA long-range synchronization reflect transient assemblies of active neuronal processing i.e. spiking in local neuronal circuits. Our results open a new avenue into measuring and understanding neuronal communication in the human brains at the level of pre-synaptic population spiking, spike synchronization, and their post-synaptic potentials in remote targets.

### Micro- and macroscale cortical architecture of HGA synchronization

HGA synchronization exhibited systematic organization at several spatial and temporal scales. At the level of cortical systems determined by fMRI intrinsic connectivity^24^, HGA synchronization was most pronounced within and between the limbic- and default-mode systems. These areas exhibit high level of activity during resting-state ^26^ and thus this finding is in line with the notion that HGA synchronization reflects neuronal processing or communication *per se*. Interestingly, in a subset of subjects who had recordings both during resting-state and during sleep, we found differences in HGA synchronization patterns between these two conditions (Suppl. Fig. 7). This suggests that HGA synchronization is state-dependent similar to fMRI resting-state connectivity in humans ^35^ and non-human primates^36^.

In addition to this insight into HGA synchronization among fMRI-derived brain systems, we examined the intrinsic community structure in the whole-brain connectome of HGA synchronization. We found evidence for significant and split-cohort reliable community structures in HGA synchronization. Together with the split-cohort reliability of the connectome itself, these findings show that HGA synchronization has a group-level stable cortical topology that is independent of individual subjects and thus also a structure that is not dictated by the individual pathogenesis or electrode placement. Moreover, this ties HGA synchronization with phase and amplitude interaction networks in slower frequencies, which as known to have salient community structures^37–40^. The HGA communities, however, were dissimilar with those at slower frequencies, which may be attributable to distinct cortical generator mechanisms of the slow and HGA signals. This conclusion was supported by the finding that at the scale of cortical laminae, HGA synchronization was significantly stronger among electrode contacts in deep than superficial layers whilst the opposite laminar organization was found for theta, alpha, and beta oscillations in SEEG here and in prior work^16, 27^. HGA synchronization thus exhibits reliable neuroanatomical organization at scales from cortical laminae to brain systems with phenomenological differences with the oscillations in slower frequencies. HGA synchronization is thus not a “by-product” of neuronal interactions coupling the slower oscillations, but rather a hitherto undiscovered component in the organization of large-scale brain dynamics.

### HGA synchronization is not attributable to artefactual sources

The connectivity and community structures of HGA synchronization as well as its laminar organization strongly suggest that it may not be attributable to physiological or technical artefacts such as signals from muscles or extra-cranial sources. Nonetheless, we corroborated this notion with a number of controls. First, in addition to white-matter referencing, HGA synchronization was observable with bipolar referencing, indicating that its current sources are millimeter-scale local in cortical tissue and very unlikely to originate from extracranial or muscular sources during inter-ictal events^41^. Second, HGA synchronization was also observable with linear-mixing insensitive interactions metric and hence not attributable to volume conduction^42^. Third, HGA synchronization was comparable among electrodes along single electrode shafts and between electrodes in different shafts, which excludes the contributions of implantation-related lesion along the shafts as well as voltage diffusion because of perfusion of CSF in the cortical lesion following shafts implant. Fourth, similar observations were made with two different SEEG data-acquisition systems and the recordings with the same electrodes in saline solution did not show any indication of artificial HGA synchronization. HGA coupling thus does not arise from the signal amplifier or data-acquisition electronics.

### HGA activity is not explained by pathological epileptic activity

The role of epileptic pathology is overall a concern for research carried out with epileptic patients. Although HGA within the epileptogenic zone show peculiar spectral and amplitude contents^43, 44^, and are temporally predictive of upcoming seizures, most recent work has questioned its pathological-only origin^45, 46^. We addressed question of physiological vs. pathophysiological genesis of HGA synchronization through several lines of analyses. First, only the putatively healthy brain areas and time-windows with no epileptic spikes were included in the primary analyses of HGA synchronization. Second, if HGA synchronization was attributable to epileptic pathophysiology, one would expect highly individual large-scale patterns, but instead of such, we observed that both the connectivity and community structures of HGA were split-cohort reliable at the group level and thus not driven by individual pathology. Third, in direct comparisons of HGA synchronization among healthy areas, between healthy areas and the epileptogenic zone, and within the epileptogenic zone, we did not find significant differences between these conditions. We found the epileptogenic zone to be distinct from healthy brain areas both in delta-frequency phase synchronization and in the phase-amplitude coupling of delta and theta oscillations with HGA. Hence, if HGA synchronization were a property of the epileptogenic zone, it would likely have been observed in this analysis. Finally, we observed no phase coupling of HGA with epileptic spikes. Taken together, HGA synchronization appears to be a property of healthy brain dynamics rather than being attributable to epileptic activity or pathophysiology.

### Putative generative mechanisms HGA synchronization

What mechanisms could underlie the signal generation of long-range phase coupled HGA? Whereas synaptic mechanisms are known to contribute to the generation of synchronized ∼200 Hz oscillations during hippocampal sharp-wave events^47^, it has remained disputed whether broad-band HGA in the 100-300 Hz range is associated with synaptic mechanisms generating neuronal population oscillations with rhythmicity in specific time scales^19^. A recent study^13^ argues that putatively-spiking-related broadband and genuinely oscillatory components with presumably synaptic-communication-based mechanisms are dissociable in the HGA signals, even though several studies suggest HGA to arise from neuronal spiking activity^48, 49^ *per se*. We speculate that the HGA signals observed in this study may reflect both local cortical population spiking activity and the consequent downstream post-synaptic potentials resulting from these volleys of spikes. Long-range MUA synchronization has not been previously reported, but it is important to note that unlike the micro-electrodes used in animal research, the larger surface area of the electrodes heavily predisposes SEEG specifically to detecting population spiking activity^50^ that is already locally synchronized in a sizeable assembly and thus able to achieve post-synaptic impact in distant targets^21^. Hence, by construction, SEEG may be effectively filtering out asynchronous multi-unit activity that is unlikely to achieve well-timed downstream effects.

## Acknowledgements

This study was supported by the Academy of Finland (project numbers: 253130, 256472, 1266745, and 266402), by the Helsinki University Research Funds, and by the Finnish Cultural Foundation (12938). The research leading to these results has received funding from the European Union Seventh Framework Programme (FP7/2007-2013) under grant agreement no. 604102 (Human Brain Project). GA was partially funded by Fondazione San Paolo (20670). The authors declare no competing financial interests.

## Data Availability

Raw data and patient details cannot be shared due to Italian governing laws as well as Ethical committee restrictions. Intermediate as well as final processed data that support the findings of this study are available from the corresponding authors upon reasonable request.

## Author contributions

AG, WSH, PJM, and PS conceived the study; AG, WSH, WN, ZA, PJM, and TB wrote the analysis code; AG, WSH, WN, HJ, TB, MMF run the analysis; CF, NL, and RA provided clinical information and preprocessed clinical data. All authors contributed to the writing and revising the manuscript. All authors read and approved the manuscript.

## Abbreviations

cPLV: complex-valued phase-locking-value (Methods eq.1)
EZ: epileptogenic zone
HFO: high-frequency oscillations (100–200 Hz)
HGA: high-gamma activity (100–300 Hz)
iPLV: the imaginary part of the complex PLV
LFP: local-field potential
nEZ: putative healthy SEEG recording sites
PLV: the absolute value of the complex PLV
SEEG: stereo-electroencephalography

## Methods

### Data acquisition

We recorded SEEG data from 67 subjects affected by drug resistant focal epilepsy and undergoing pre-surgical clinical assessment for the ablation of the epileptic focus. We acquired monopolar (with shared reference in the white matter far from the putative epileptic zone) local field potentials (LFPs) from brain tissue with platinum–iridium, multi-lead electrodes. Each penetrating shaft has 8 to 15 contacts, and the contacts were 2 mm long, 0.8 mm thick and had an inter-contact border-to-border distance of 1.5 mm (DIXI medical, Besancon, France). The anatomical positions and amounts of electrodes varied according to surgical requirements ^1^. On average, each subject had 17 ± 3 (mean ± standard deviation) shafts (range 9-23) with a total of 153 ± 20 electrode contacts (range 122-184, left hemisphere: 66 ± 54, right hemisphere: 47 ± 55 contacts, grey-matter contacts: 110±25). We acquired an average of 10 minutes of uninterrupted spontaneous activity with eyes closed in these patients with a 192-channel SEEG amplifier system (NIHON-KOHDEN NEUROFAX-110) at a sampling rate of 1 kHz. Before electrode implantation, the subjects gave written informed consent for participation in research studies and for publication of their data. This study was approved by the ethical committee (ID 939) of the Niguarda “Ca’ Granda” Hospital, Milan, and was performed according to the Declaration of Helsinki.

### Signal pre-processing

We excluded electrode contacts (1.3±1.2, range 0–10, contacts) that demonstrate non-physiological activity from analyses. We employed a novel referencing scheme for SEEG data where electrodes in grey-matter were referenced by the contacts located in the closest white-matter (CW)^2^. This referencing scheme is proven optimal for preserving phase relationship between SEEG contact data^2^. The final size of channels analysed is on average 110±25 for each subject and 7491 in total.

Prior to the main analysis, SEEG time series were filtered with 18 Finite Impulse Response (FIR) filters (equiripples 1 % of maximal band-pass ripples) with central frequency (F_c_) ranging from 2.50 to 320Hz. We used a relative bandwidth approach for filter banks such that pass band (W_p_) and band stop (W_s_) were defined as 0.5×F_c_ and 2×F_c_, respectively for Low- and High-pass filters separately. We excluded all 50 Hz line-noise harmonics using a band-stop equiripple FIR filter with 1 % of maximal band-pass ripples and 3 up to 8Hz width for the stop band parameters.

Epileptic events such as interictal spikes are characterized by high-amplitude fast temporal dynamics as well as by widespread spatial diffusion. Due to possible filtering artefacts around epileptic spikes and the resultant increase in synchrony, we discarded periods of 500ms containing Interictal Epileptic Events (IIE). We defined such periods as the temporal windows where at least 10% of cortical contacts demonstrated abnormal concurrent sharp peaks in more than half of the 18 frequency bands. Such episodes were excluded from within- and cross-frequency synchrony analysis. To identify such periods in each contact, we partitioned the contact amplitude envelope time series into 500 ms non-overlapping windows and marked IIE events as the time windows within which at least 3 consecutive samples are 5 times the standard deviation above the mean amplitude.

### Defining the epileptic zones based on seizure activities in SEEG signals

The epileptogenic and seizure propagation zone were identified by clinical experts by visual analysis of the SEEG traces^1, 3^. *Epileptogenic* areas are the hypothetical brain areas that are necessary and sufficient for the origin and early organization of the epileptic activities ^4^, from where contacts recording often show low voltage fast discharge or spike and wave events at seizure onset. Seizure *propagation* area are recruited during the seizure evolution, but they do not generate seizures ^5, 6^, from where contact recording show delayed, rhythmic modifications after seizure initiation in the epileptogenic areas. In this study, we combined epileptogenic and propagation areas as the epileptogenic zone (EZ) to distinguish from the rest of brain areas that are referred to as *putative* healthy zones (nEZ).

### Functional connectivity estimates

We estimated inter-areal phase-phase interactions at individual subject level using the Phase Locking Value (PLV). Defining *x*′(*t*) = *x*(*t*) + *i*H[*x*(*t*)] as the analytical representation of the signal *x*(*t*), where H[··] denotes the Hilbert transform, complex PLV (cPLV) is computed as ^7^:

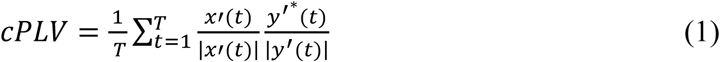

where *T* is the sample number of the entire signal (*i.e.*, ∼10 minutes), and * is complex conjugate. We computed cPLV for the entire recording excluding 500 ms time windows showing epileptic or artefactual spikes (see below). The PLV is the absolute value of complex cPLV (*PLV* = |*cPLV*|), and it is a scalar measure bounded between 0 and 1 indicating absence of phase and full phase synchronization, respectively.

Additionally, we used imaginary part of cPLV (*iPLV* = *Im(cPLV)*), a metric insensitive to zero-lag interactions caused by volume conduction ^8–10^, for verification. For both PLV and iPLV connectivity, the fraction of significant edges (*K*) is the number of significant edges divided by the total possible edge number. Since one same white-matter contact can be used for referencing multiple cortical contacts, we rejected derivations with shared reference.

### Statistical hypothesis tests

We estimated the null-hypothesis distributions of interaction metrics with surrogates that preserve the temporal autocorrelation structure of the original signals while abolishing correlations between two contacts. For each contact pair, we divided each narrow band time series into two blocks with a random time point *k* so that *x*_1_(*t*) = *x*(1 … *k*) and *x*_2_(*t*) = *x*(*k* … *T*), and constructed the surrogate as *x*_*surr*_(*t*) = [*x*_2_, *x*_1_]. We computed surrogate PLV across all channel pairs and assembled the surrogate interaction matrix, and its mean and standard deviation was later used in hypothesis testing.

### Correlation estimates post-processing

To demonstrate how interaction strength varies as a function of spatial distance between recording sites, we divided the inter-contact distances into three ranges (short (SH) 2 cm ≤ *x* < 4.6 cm; medium (MD) 4.6 cm ≤ *x* < 6 cm, and long-range (LG) 6 ≤ *x* < 13 cm) with same number of edges falling into each distance range. Therefore, we averaged across subjects all the edges falling within each distance range and for each frequency band separately (N=48702/range). The confidence intervals for PLV and iPLV, were expressed relatively to the surrogate means (SM) for PLV (3.42*SM corresponding to *p* < 0.001, Rayleigh distribution), and the surrogate standard deviations (SD) for iPLV (3.58*SD corresponding to *p* < 0.001, normal distribution).

To compare signals from superficial and deep layers in the grey matter (Fig. 4 and Suppl. Fig. 3), we divided contacts into “shallow” and “deep” based their Grey Matter Proximity Index (GMPI)^2^ that is defined as the relative distance between the contact location and the nearest white-grey border surface, normalized by the grey matter thickness at that location:

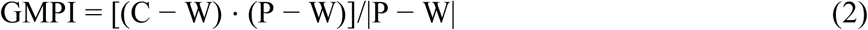

where P(x, y, z), W(x, y, z) and C(x, y, z) are the vertices on the pial, white-matter surface and contact coordinates in 3D individually reconstructed brain from MRI scan, respectively. Values 0 < GMPI < 1 indicate that the contact midpoint is located in grey matter whereas a negative GMPI indicates that the contact midpoint is in the white matter.

We set the criteria −0.3 < GMPI < 0 and 0.5 < GMPI < 1.2 to classify deep and superficial layer contacts respectively. Next, PLV and iPLV estimates were averaged across subjects between deep-deep (DD) and superficial-superficial (SS) contact-pairs. We tested for between groups difference with a paired permutation test (100 random samples created by shuffling SS and DD labels within subjects; threshold for significance corrected for multiple comparisons with Bonferroni *p* < 0.05/N, with N=18).

### Anatomical co-localization of SEEG implant and Functional System characterization

To show that high-gamma phase coupling is associated with a common systems-level mechanism, we assessed the fraction of significant edge (*K*) between functional systems^11^. We extracted cortical parcels from pre-surgical T1 MRI 3D-FFE (used for surgical planning) using Freesurfer ^12^, and we used the new Schaefer parcellation ^13^ that favours functional networks topology over structural (gyral) topology ^14^. The resulting cortical meshes were divided into seven functional systems: Visual, somato-motor (SM), Dorsal Attentional (DAN), Ventral Attentional (VAN), Limbic, Fronto-parietal (FP), and Default Mode Network (DMN). This atlas does not include subcortical regions and thus subcortical contacts were discarded from this analysis. We thus assigned each cortical contact to a cortical parcel that belongs to a functional system. We then computed the fraction of significant edges (*K*) between 7 functional systems.

Note that we initially conducted this analysis in the Destrieux 148-parcel atlas ^14^, but we re-analysed the whole dataset with the Schaefer atlas ^13^ after its release for both verifying the observations in the Destrieux atlas and for achieving optimal parcel-to-system morphing quality.

### Cross-frequency coupling of slow rhythm phase and fast rhythm amplitude

Two signals of distinct rhythms are cross-frequency phase-amplitude coupled (PAC) if the phase of a slow neuronal oscillation modulates the amplitude fluctuations of the faster neuronal oscillations. PAC can be estimated using phase synchronization, Euler’s formula, or examining whether the power of fast rhythms is non-uniformly distributed over low-frequency phase ^15–19^.

The rationale is that if the power fluctuations of fast rhythms are modulated by the phase of the slow oscillations, the fluctuations of these two time-series should be synchronized. In this study, we estimated PAC with the phase locking value (PLV) as:

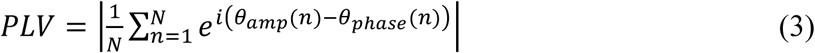

where *θ_amp_*(*η*) is the phase time series of the power envelope of fast rhythm while *θ_phase_* (*η*) is the narrow band phase time series of the slow rhythm. When there is a consistent relationship between these two time-series, the vector length of the mean phase differences (in the polar coordinate across all *n* samples) should be greater than zero, and a maximum value of 1 indicates perfect coupling. The significance of PAC PLV value was determined in the same manner in individual subjects as we conducted for 1:1 phase synchrony described earlier.

### Detecting the modular structures

If observed individual level long-range phase synchrony between SEEG contacts are truthful, we next ask ourselves whether, on the population level, the observed synchrony networks are functionally meaningful; in other words, whether the networks can be reliably subdivided into well-separated functional subsets of nodes, or ‘modules’, across frequencies and in particular in the HGA band.

To answer this question, we first unambiguously assigned each cortical SEEG contact to one of the 100 Schaefer parcels^13^ (subcortical and EZ cortical contacts were not analyzed). For each frequency, we pooled PLV values of contact-pairs originating from homologous parcel-pairs across population, and then assigned to each parcel-pair the median of these pooled PLV values, defining this median value as the weight of the edge between the selected parcels. The median was used because of the non-normality of PLV distribution. Finally, we removed those edges that had not been sampled at least 20 times in total, and by at least 3 different subjects. By doing so, we collapsed sparsely sampled, patient-specific contact-pair networks into well-sampled, population-level phase synchrony networks, defined over brain regions (see Fig. 1c). Due to the low inter-hemispheric contact coverage, we limited our investigation to intra-hemispheric modular structures.

Given the sparse and non-homogenous sampling of cortical space across patients, 10% to 20% of all possible parcel-pairs have not been sampled, *i.e.*, matrices have missing values (Fig. 1c). We assume that the missing PLV values come from the same probability distribution of the observed part of the corresponding network. Under this assumption, for each frequency, we generated 1,000 variants of the original phase synchrony network, by filling in each missing edge with the PLV value of an existing edge, randomly and independently selected. We then used a consensus clustering approach^20^ over these 1,000 variants for the computation of the modules. Thereby, the modules successively obtained from these ‘filled’ networks are less likely confounded by distributed local fragmentation of the network topology due to missing values.

We applied the Leiden algorithm^21^ to identify modules in each of the 1,000 variants. The resolution γ parameter of the method weighs the importance of the null model (i.e., random network with no modular structure) against which the original network is compared, when identifying the network partition which maximise the modularity value:

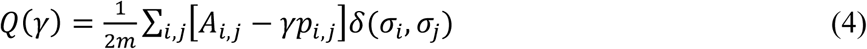

where *Q*(*γ*) is the modularity value at resolution parameter *γ*, *m* is the total strength of the network, *A*_*i*,*j*_ is the network value in row *i* and column *j*, while *p_i,j_* are the network values in the null model, i.e. expected by chance for row *i* and column *j*. Here, 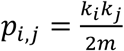, where *k*_*i*_ and *k*_*j*_ are the total strength of network values in row *i* and column *j* respectively; *δ*(*σ*_*i*_, *σ*_*j*_) is equal to 1 when node *i* and node *j* belong to same community, and 0 otherwise. The modularity value measures the quality of the network partition: a good partition should maximize the connections between nodes in the same module, while minimizing the connections between nodes in different modules. Values of γ less than 1 tend to favour a small number of large communities, since the null model is down-weighed; on the contrary, values of γ larger than 1 tend to produce a high number of small communities, since the importance of the null model is up-weighted. We computed module partitions for a range of values for γ, from 1 to 1.5, with steps of 0.05.

For each frequency and value of γ, we computed a single partition from the 1,000 variants of the original network, using a consensus approach^20^. Briefly, we first computed a modular partition for each of the 1,000 variants of the network; from this partition we derived a binary community co-assignment matrix *C*, where *C_ij_* equals 1 when brain region *i* and *j* are assigned to the same module and 0 otherwise. This was done in order to be able to compare and pool together different partitions, despite possible differences in the numberings of the modules across partitions. We then averaged the 1,000 community co-assignment matrices into a single weighted matrix, where each weight estimates how often the corresponding pair of nodes were assigned to the same module. We fed this final matrix again to the Leiden method, with the same γ value used on the 1,000 variants, to produce a single final partition for the given γ. This consensus clustering method considers the set of 1,000 variants of the functional network as ‘noisy’ versions of the true underlying network: by averaging their partition assignments, the graph noise at individual level can be mitigated.

We also evaluated the similarity between two partitions *m* and *n* across frequencies and resolutions. Here, *m* and *n* are two vectors with as many elements as the cortical parcels: the value of each element is the ID of the module to which the parcel was assigned in the partition. For each *m* and *n*, we created the community co-assignment matrices *C*^(*m*)^ and *C*^(*η*)^ as described earlier, and computed the partition similarity between *m* and *n* as^22^:

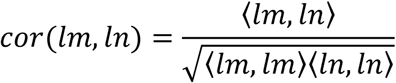

Where 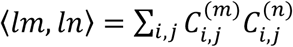. Since the dot product 〈*lm*, *ln*〉 satisfies the Cauchy-Schwartz inequality, such that 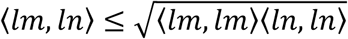, the partition similarity equals 0 for uncorrelated partitions and 1 for identical partitions. We only computed the similarity between partitions of the same hemisphere of the brain, and not between partitions of different hemispheres because, in the Schaefer atlas, the subdivision into brain region is asymmetric across hemispheres, and the number and demarcation of nodes is different.

Lastly, we assessed the range of resolution parameter values at which modules could be reliably identified. This was done both at the level of the entire network as well as at the level of individual regions (see Suppl. text).

### PLV in amplitude bins

To assess whether larger values of phase synchrony were correlated with higher amplitude values, we estimated instantaneous amplitude and phase profiles of the filtered time-series by means of Hilbert transform. We then normalised each amplitude time-course by its median and divided the normalized amplitude samples in quintiles. For each frequency, we built an amplitude-amplitude correlation matrix containing the instantaneous phase difference between channel pairs of each time-samples at that amplitude bins. We discarded amplitude samples larger than twice the amplitude median to remove effects of subthreshold spikes. We quantified the number of time-samples falling in each bin as a simple amplitude correlation measure. We hypothesized that that, if a given contact pair is amplitude correlated, the time-samples would not be randomly distributed over amplitude bins. Indeed, it will result in highly skewed distribution of time-samples towards larger amplitude bins. On the other hand, in the absence of real amplitude correlation that distribution would be undistinguishable from a uniform distribution. Hence, to test for a moment-to-moment amplitude correlation, we quantified the distance of the time-sample distribution from a uniform distribution for each amplitude-amplitude bin under the above null-hypotheses of no correlation. To quantify whether phase consistency was correlated with moment-to-moment amplitude modulation, we quantified PLV in each amplitude-amplitude bin. The PLV is a measure sensitive to the sample number used^23^, hence we quantified the minimum number of samples falling in each bin and then quantified PLV in amplitude bin with matched time-samples. Specifically, for each contact pair, we computed instantaneous phase difference across the entire time course. By grouping instantaneous amplitude samples falling in same bin, we averaged phase differences in each amplitude bin and later averaged this quantity across channel pairs and subjects.

### Evoked Phase Synchronization

In order to investigate whether HG synchrony was functionally relevant in cognitive tasks, we recruited 11 subjects and had them perform a visuo-motor Go/no-Go task^24^. Subjects were asked to respond as quickly as possible to a Go cue (blue rectangle) by pressing the space bar on a standard keyboard and ignore the no-Go cue (yellow rectangle). After an initial two minutes of resting state (used as control condition in the following analyses), subjects performed a total of 1000 trials (750 Go trials). Pre-processing of SEEG data remained unchanged compared to resting-state analyses. Specifically, we removed any defective contacts as well as those recording from subcortical structures. We used CW referencing as reference schema and removed all common-reference contact pair for further pairwise analyses. Filter settings and spectral resolution remained unchanged as in the resting-state case described above. Three patients showed no responses in any channels and thus were excluded from further analyses.

To assess whether a cortical response could be measurable after stimulus onset, we measured Event Related Synchronization and Desynchronization (ERSD) for each frequency separately. For each CW-referenced SEEG contact, we averaged responses across trials, then normalised evoked responses by subtracting the mean baseline amplitude (from 500 to 10 ms before stimulus onset) and smoothed with a moving average FIR filter of 25 ms length in order to more reliably estimate peaks in amplitude response. Finally, we normalised (z-score) each ERSD and, for each subject, we divided the contacts in two groups (task-relevant and -irrelevant) of 30 channels each based on their average amplitude response during 200–550 ms after NoGo cue onsets.

We next compared phase synchronization between resting and task conditions in these eight subjects. We computed the cPLV dividing the task data (on average 15 minutes) in 2 minutes (*i.e.,* equivalent to the duration of initial resting period) long non-overlapping windows in order to account for the sample-size bias of PLV estimates under the null-hypothesis of no coupling, *i.e.*, the larger the sample size the smaller the surrogate PLV and hence the more significant the effect might appear. For the evoked phase synchrony, we divided each trial (–500 ms to 800 ms around visual cue onset) in thirteen 100 ms long windows. Then, we computed the time-resolved cPLV in each time window by averaging phase differences within each time windows across trials. Evoked cPLV surrogates were constructed by randomly shuffling trials of one channel while keeping unchanged the trial order of the other one. We also replicated our observation of time-resolved cPLV for five 260 ms long windows (data not shown).

### Statistical assessment of task relevant effect

We first performed a permutation test for rest vs task condition on static iPLV. Hence, we pooled phase synchrony estimates across subjects and randomly permuted 10,000 times rest and task labels. Then, we computed the *p*-value as the fraction of permuted elements whose iPLV rest-task difference exceeds the observed (non-permuted) difference using the exact *p*-value approximation described in^25^.

Finally, to describe effects of cognitive load on phase synchronization compared to surrogates and between task-relevant and irrelevant channels, we performed a randomization max T test with *p* < 0.05. In this test for each comparison, we picked the maximum value of test statistics (iPLV difference) across time-bins for all 1000 permutations. We then computed the *p* < 0.05 confidence limit on these 1000 permuted max T and tested whether the observed iPLV difference was above that threshold in any time bins for both surrogates and task-irrelevant/relevant tests.

## Supplementary Text

### Is high-gamma synchronization attributable to artificial sources?

#### Cohort demographics and sampling statistics

The majority of the patients had SEEG electrodes implanted either in left or right hemisphere, and 11 subjects had bilateral implantation. There was some variability in non-epileptic cortical contact number per patient (**b**) and distinct patients per cortical parcel (**c**), but contact number and fraction of subject per parcel was correlated (r= 0.9), thus ruling out the possibility that a subset of subjects is biasing the connectivity estimates on parcel-to-parcel scale. On the systems-level the right hemisphere was sampled more extensively than the left hemisphere (d), but note that the majority of the left hemisphere functional systems was sample with more than 20 distinct subjects. Therefore, the connectivity is well sampled on the systems-level as well.

For the majority of the connectivity analyses, we included only the tentative healthy contacts (Fig 1, Suppl. Fig. 1 c,d), but we also analyzed the epileptogenic (EZ) contacts for some comparisons. The spatial sampling of the EZ contacts is shown in panels **e** and **f** of Suppl. Fig. 1.

### HGA phase synchrony is a property of inter-areal coupling in human brain and is not confounded by specific referencing scheme or volume-conduction

#### Referencing scheme

Although local white matter referencing effectively suppresses volume-conducted signal components and best preserve phase dynamics^1^, we employed two extra control analyses to confirm that high-gamma synchronization was not attributable to referencing scheme and/or volume conduction.

First, we inspected the mean PLV as a function of frequency using the classical bipolar referencing (*i.e.*, each contact is referenced by its neighbouring contact). With a center-to-center separation of 3 mm between two neighbouring contacts, bipolar signals reflect strictly local neuronal activities. However, it can also distort signals due to cancellation, especially when neighbouring contacts are located in functionally/anatomically different regions ^1^. The mean PLV and the fraction of significant PLV of bipolar data (Suppl. Fig. 1g) had no visible differences from corresponding results with white matter referencing (Fig. 2a, b).

#### Volume conduction

The PLV is known to be contaminated with spurious correlations due to volume conduction or field spread in macro-scale, on-scalp measurement of the cortical activity such as EEG and MEG. We excluded nearby contacts from phase synchrony analysis, but to further prove the observed HGA synchrony with PLV was not due to volume conduction, we inspected the phase synchronization using the imaginary part of the complex-valued PLV (iPLV). The iPLV is insensitive to instantaneous linear signal mixing effects ^2–4^. However I did not use it as a main connectivity metric because it does not report true near zero-phase-lag interactions, and its ambiguity: a change in the iPLV value can be caused by a change in phase locking, a change in the phase lag, or the combination of both.

We found that the HGA synchrony remained prominent when estimated with iPLV for both closest-white-matter (Suppl. Fig. 1h) and bipolar (Suppl. Fig. 1g) referencing. Hence, a significant amount of the long-range HGA synchrony emerged between local populations have non-zero phase lag (*i.e.*, iPLV remove phase correlations with near zero phase-lag).

Last, with bipolar referencing, we reproduced similar results on synchronization profiles between cortical layers (Suppl. Fig. 2a, see Fig 4). Bipolar approach mixes neuronal sources coming from superficial and deep layers given the contact edge-to-edge distance of 1.5 mm and the contact dimensions of 2 mm which are comparable to the average cortical thickness in adult brain 4mm^1^. Interestingly, phase synchrony was not different between deep and shallow bipolar referenced contacts (permutation test, *p* < 0.05 Bonferroni corrected with N=18) for slower rhythms (< 100Hz) (Suppl. Fig. 2a). This might arise from the fact that slower oscillations have been demonstrated to spread much further in distance compared to HGA, hence leading to a cancellation between layers due to the inappropriate bipolar referencing (*i.e.*, the main merits of the closest-white-matter over bipolar referencing). On the other hand, the highly local nature of HGA still yields significant difference between shallow and deep contacts even at long-range distances. This suggests that HGA synchrony is an actual property of the human cerebral cortex and that such synchrony is tightly related to cortical laminar architecture.

In summary, the parallel set of analyses conducted using bipolar montage and iPLV (Suppl. Fig. 2, 3) effectively refuted that the HGA phase coupling were attributable to extra-cranial sources of physiological artefacts, such as eye or scalp muscle activities, or external device related noise patterns. Such sources of non-neuronal origin could not produce electric field gradients steep enough to be observable in bipolar recordings while maintaining systematic long-range phase relationships with a non-zero-lag between widely separate and highly local cortical SEEG signals.

### Line-noise leakage does not confound HGA phase synchrony

To test whether an insufficient attenuation in band-stop filters left residual artefactual power from line-noise sources, we divided subjects in two groups based on relative quantity of line noise power in respect to side bands (see Methods). Properly controlled setup has already ruled out much contribution from line-noise: 6 subjects showed peaks in power spectrum profiles (Suppl. Fig. 3a). We used k-means algorithm to split population in two groups using a 9 dimensional parameter-space described by the relative power of line-noise central peak, *i.e.* 50Hz, and its 6 harmonics (100, 150, 200, 250, 300, 350, 400, 450 Hz) in respect to the side bands (5Hz each). Principal Component Analysis (PCA) suggested that two groups could be identified (Suppl. Fig 3b, their power spectrum see c, d). Finally, we computed mean PLV and iPLV (Suppl. Fig. 3e, 3f) for both groups and found no differences between groups. Thus, we concluded that line-noise does not confound observed HGA phase synchronization.

### Split-cohort reliability of strength, density and connection pattern of HGA synchrony

The HGA synchronisation had split-cohort reliability in PLV strength and *K* on contacts level and systems-level (Suppl. Fig. 4 a, b & d, e, see Fig. 2a, b) and pattern of functional connections (Suppl. Fig. 4c, right). We split the subjects into two cohorts (34, 33 subjects respectively) so that match between number of contacts per region in the two cohorts was maximised (Suppl. Fig 4c, left). It is inconceivable that the across-subject reproducibility of large-scale cortical architecture could arise from technical artefacts or extra-cranial sources in a cohort where each subject has a unique pattern of SEEG shaft implantation.

### Spectral leakage from lower-frequency processes and filtering approach

We considered whether high-gamma synchronization could be explained by inadequate attenuation in band-pass filtering, which would cause leakage of low-frequency components into the high-gamma band. This confounder was ruled out due to absence of any low-frequency components in the filtered time series (Suppl. Fig. 5a). From the physiological perspective, the dissimilarity between the community structures high-gamma synchronization to reflect a facet of neuronal activity that is distinct from the lower-frequency (< 100 Hz) signals.

Moreover, to exclude the possibility that the observations of HGA could be biased by the finite impulse response (FIR) filtering approach, we used Morlet filtering to reproduce all principal analyses. We found Morlet wavelets (Suppl. Fig. 5b) to yield similar results as FIR filters (see Fig. 2a, b) for closest-white-matter referenced data.

### Ruling out other possible confounders: Dissociation of oscillatory from unitary spike-like generators of high-gamma activity

#### Technical confounder from the amplifier

Given the relatively small amplitudes of cortical oscillations in high-gamma range, we asked whether observed phase synchrony could possibly be due to synched noise generated within amplifiers. At the beginning of 2017 the Niguarda Hospital renewed the acquisition system to a more recent version from Nihon-Kohden. Hence, we acquired 10 new subjects (data not included in main results but presented here) as well as 10 minutes data from two electrodes (18 contacts each) immersed in a saline solution. We then performed the same analysis pipeline on these data and observed that saline solution yields no synchrony in any frequency range (Suppl. Fig. 6a). Furthermore, the strength of between-contact phase synchrony from saline solution test is close to the surrogate of patients data but smaller than the HGA synchrony strength. Thus we concluded that the observed HGA synchrony is not due to artificial sources from the acquisition system.

#### Spikes are not correlated with synchrony strengths

Theoretically, transient spike-like neuronal events could be picked up by filtering in the high-gamma frequency band. We effectively discarded all time-windows containing epileptic spikes (referred to as “cleaned-data”, see main text and Methods) that could possibly introduce spurious correlations. We found that the number of spikes detected were not correlated with the mean PLV across most of the frequencies except for 2.5 Hz (Suppl. Fig. 6c), which rules out the phase synchronization observed in the gamma to ripple frequency bands is artificial due to the epileptic or inter-ictal spikes.

#### No evidence of muscular activity contributions to HGA synchrony

To further assess whether muscular activity inflated the PLV estimates, we hypothesized that contacts closer to skull would have picked more muscular activity compared to contacts recording far from it. Hence, we analyzed separately contacts recording far-from-skull and those near-skull. We observed that former type of contacts show increased phase synchrony compared to latter (Suppl. Fig. 6d) further suggesting absence of confounds from muscular artefacts.

However, sub-thresholds spikes could still artificially inflate phase synchrony. We localized all high-gamma amplitude peaks from our cleaned-data and assessed the phase distributions of high-gamma oscillations at the amplitude peaks for channel pairs exhibiting significant (*p* < 0.001) long-range phase coupling. If spikes played any role in the generation of these signals, the phase distributions would have a peak at zero-lag whereas for ongoing oscillations, these distributions are indistinguishable from a uniform distribution. We set the amplitude peak detection threshold to mean plus three times the standard deviations of the filtered amplitude time series, which yielded 1523.06 ± 231850 (mean ± SD) peaks per electrode contact.

We found that 81.2% ± 0.29% of channels show phase distribution at amplitude peaks indistinguishable (*p* > 0.001, Rayleigh test for uniformity) from uniform phase distribution. This observation further consolidates the notion of these high-gamma observations are reflecting genuine neuronal oscillations.

### The synchrony was different between sleep and wakeful resting

To further support physiological relevance of the HGA synchrony, we measured phase synchrony with iPLV during slow-wave sleep (SWS) in 7 subjects and compared with iPLV during resting state for the same 7 subjects. The matrices of mean iPLV (across HGA frequencies) between functional systems were different between rest and sleep conditions (dissimilarity permutation test, *p* < 0.001, N = 1,000, one-tailed), (Supp. Fig. 7a,b). Distance between rest and sleep mean PLV matrices was quantified using dissimilarity, *i.e.*, 1-correlation, between the matrices. Null distribution of dissimilarity was estimated by computing dissimilarity between pairs of surrogate rest and sleep matrices, generated by random mixing (without replacement) 1,000 times of the rest and sleep subject-level matrices prior to estimating the surrogate group-level rest and sleep matrices. Furthermore, higher phase synchrony, as measured with iPLV, was found for slow-wave sleep within the limbic system and between the limbic and default-mode and ventral-attention systems (two-tailed t-test, p < 0.05, uncorrected).

These findings together with all above proves that high-gamma synchrony is a general property of cortical dynamics, which is modulated by altered conscious states during SWS and hence cannot be attributable to artefactual source.

### Additional PLV in AA bins frequency

To further corroborate that HGA synchrony is mostly correlated with HG amplitude bursts rather than with background activity, we showed that higher PLV values are measured in correspondence with concurrent amplitude bursts between contacts. This main effect is shown in Fig.5, and here we show the other ranges of HGA (Suppl. Fig. 8).

### Community structures are stable at a range of resolution parameter values

To assess the range of resolution parameters at which communities could be identified, we performed tests at the level of individual regions and also at the level of the whole network. The range of resolution parameter values explored was 1 to 1.5 and for this range, the number of modules identified was between 2 and 13 (Suppl. Fig. 9a).

At the level of the whole network, we determined if the ‘modularity’ value of the network, *i.e.* the extent to which it can be divided into non-overlapping communities, was higher than that of equivalent random networks. We found that across the high-gamma frequencies, there was a range of resolution parameters for which the networks were significantly modular (permutation test, *p*<0.05, N=100, one-tailed) (Suppl. Fig. 9b). This was so for both left and right hemispheres.

To determine the range of resolution parameters at which significant modular structure can be detected, we first identified modules at γ values from 1 to 1.5 using consensus clustering with 100 repetitions (see Methods). For the identified modular structure at each γ value, the ‘modularity’ output parameter was obtained, *i.e.*, the extent to which network can be sub-divided into non-overlapping modules. To determine if this modularity value was statistically significant, we generated equivalent random networks of the original network, by randomly rewiring the edges while maintaining degree and strength distributions of the original network. Modules were then identified for this ensemble of 100 equivalent random networks, for the gamma values between 1 and 1.5, and the corresponding modularity value was obtained. This procedure was repeated 100 times, to obtain a distribution of modularity values corresponding to equivalent random networks. For the entire set of gamma values, modularity from the original network was z-scored with respect to the distribution of modularity values for equivalent random networks. The γ values for which modular structure had a z-scored modularity value > 2 were marked as statistically significant. This was done for frequencies from 113 Hz to 320 Hz, and for two hemispheres separately.

At the level of individual regions, we assessed the percentage of stable regions, *i.e.* the percentage of regions for which modules could be assigned reliably (Suppl. Fig. 9c). For each of the high-gamma frequencies, across the range of resolution parameters studied, we found a statistically significant percentage of stable regions (bootstrapping test, *p*<0.05, N=100, one-tailed) for both left and right hemispheres.

To determine the reliability with which each brain region can be assigned to its module, we generated a set of 100 functional connectomes for each HGA frequency band by bootstrapping (with replacement) across subjects. For 1< γ < 1.5, we identified modules for each of the bootstrapped connectomes, via consensus clustering with 100 repetitions (see Methods). Then, for each brain region, we generated a vector indicating the set of regions in its same module (indicated by 1) and those regions in other modules (indicated by 0). This vector was compared to the corresponding vector from the modular structure of the original functional connectome. In particular, we estimated the sum of the number of regions that were correctly located in the same module and the number that was correctly located in other modules (compared to the original modular structure), as a function of the total number of regions (excluding own region). This was done for each of the 100 bootstrapped functional connectomes and gave a value between 0 and 1, quantifying the similarity of the modular structure between the original and bootstrapped network, for that region. Values close to 1 across the 100 bootstrapped networks indicate that the assignment of the region to its module in the original network is reliable. To ascertain a statistically significant and reliable module assignment, we created a null-distribution by estimating the same measure when the vector indicating the modular structure of a region is randomly permuted 100 times (without replacement). The module affiliation of a region was then considered to be stable if the mean ‘reliability’ of the region across the 100 bootstrapped networks was higher than the 95^th^ percentile value of the corresponding null distribution, i.e. mean ‘null reliability’ across permutations, for the 100 bootstrapped networks. The percentage of stable regions was considered statistically significant when it was higher than 5% of the total number of regions, which was the expected false positive rate.

---

**Figure S1.**
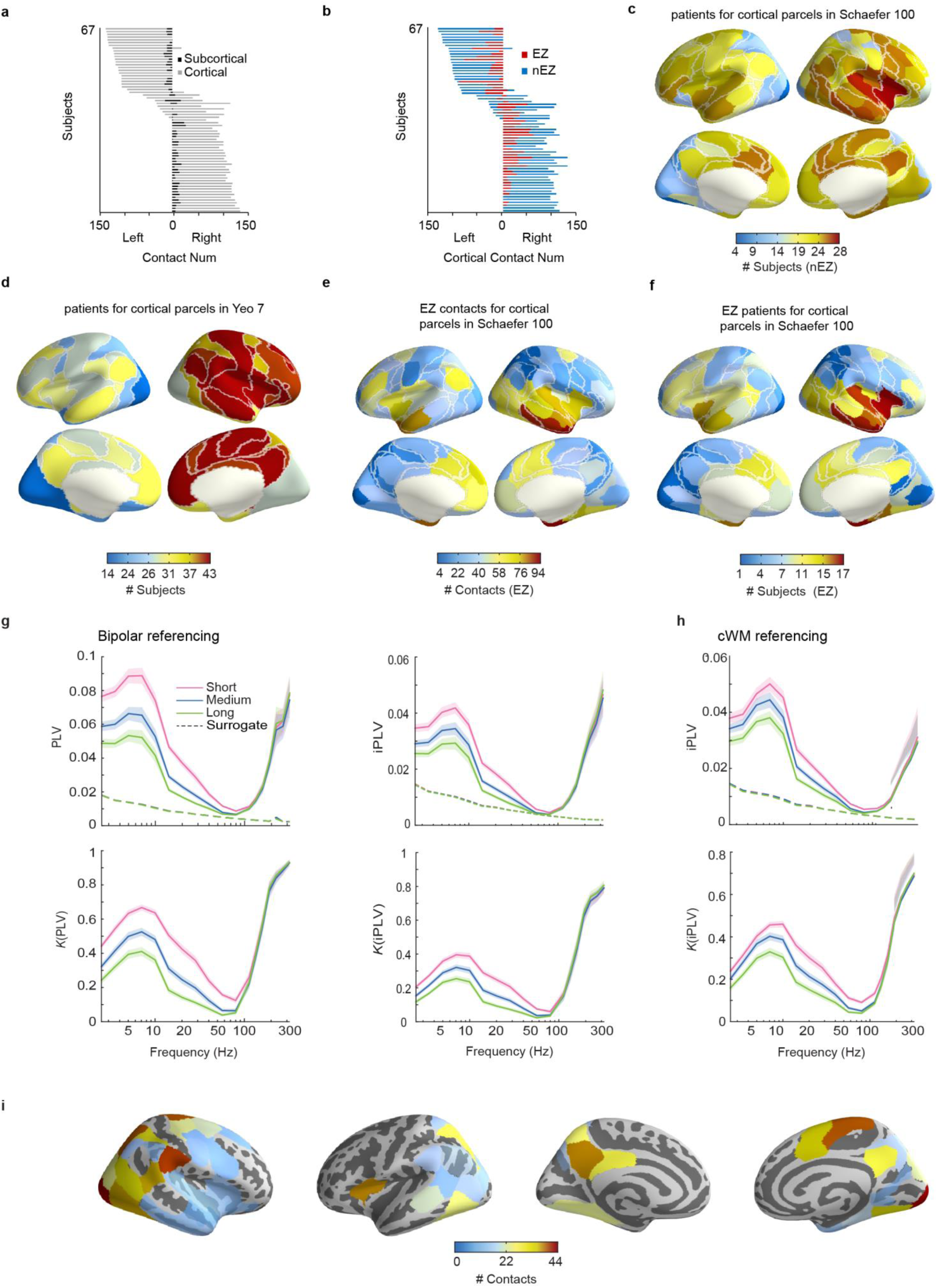
Cohort demographics and supporting evidences against confounds by specific referencing scheme or volume conduction. **a,** Number of SEEG electrode contacts per subject in cortical and sub-cortical regions (in left or right hemisphere). **b,** Number of epileptic (EZ) and healthy (nEZ) cortical contacts. **c,** Number of subjects sampled in each Schaefer cortical parcel using nEZ contacts only. **d,** Number of subjects sampled in each of the 7 functional systems. **e,** Number of distinct EZ contacts across subjects for each of 100 cortical parcels (Schaefer et al, 2017). **f,** Number of subjects with at least one EZ contact in each of 100 cortical parcels (Schaefer et al, 2017). **g,** The main finding of HGA phase synchrony (see Fig. 2) persists when using the *bipolar* referencing scheme and when synchrony was estimated with iPLV and *K* of iPLV; shaded areas represent confidence limits (two-tail; *p* < 0.05) for bootstrapped values (N = 100). Dashed lines represent surrogate (N = 100) data level for *p* < 0.001. **h,** iPLV and connection density *K* for closest-white referenced data. **i,** Number of cortical contacts in the Go/NoGo cohort.

**Figure S2.**
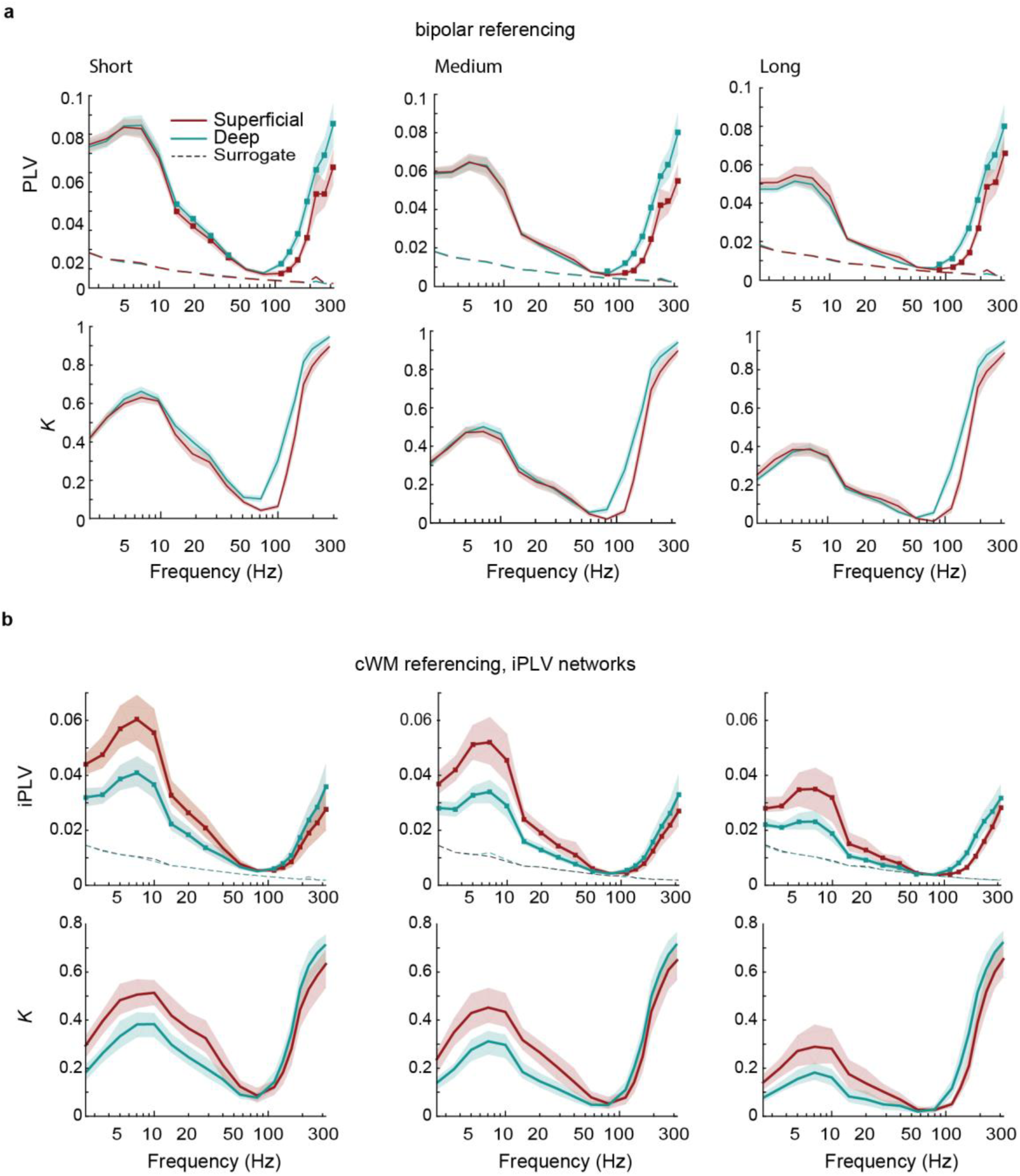
High-gamma phase synchrony also differs between cortical layers in bipolar-referenced data. **a,** Different layer profiles in HGA frequencies for bipolar-referenced data at short, medium and long distances. Shaded areas represent 2.5% and 97.5% confidence limits for bootstrapped values (N = 100). Square markers represent significance for a two-tail permutation test (N = 100) over edges with *p* < 0.05. **b,** Mean phase synchrony as estimated with iPLV and *K* of iPLV of closest-white-matter referenced data also shows distinct layer profiles.

**Figure S3.**
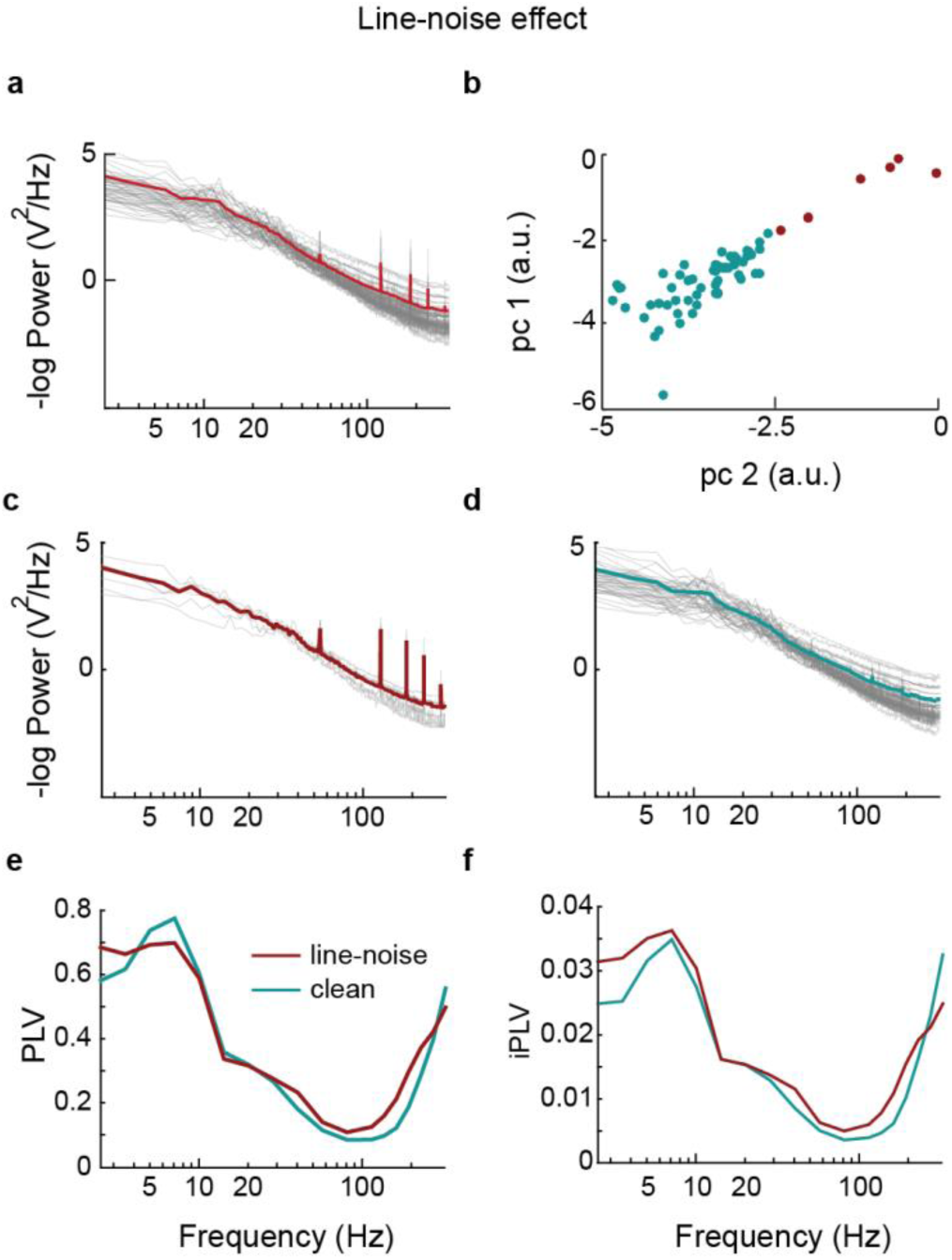
Line noise harmonics are not correlated with increased phase synchrony. **a,** Group average (red) and single-subject (gray) mean power (across channels) of the unfiltered raw time-series show the presence of line-noise peaks. **b,** PCA-decomposed power ratio of central line noise harmonics relative to side-flanks show a clear separation between affected (red) and un-affected (azure) subjects. **c-d,** Within-group mean power for unaffected (azure) and affected (red) subjects show that the vast majority (61/67) of data are not affected by line noise. **e-f,** No prominent differences in mean PLV and mean iPLV are observed between the two groups, indicating complete suppression of line-noise artefacts.

**Figure S4.**
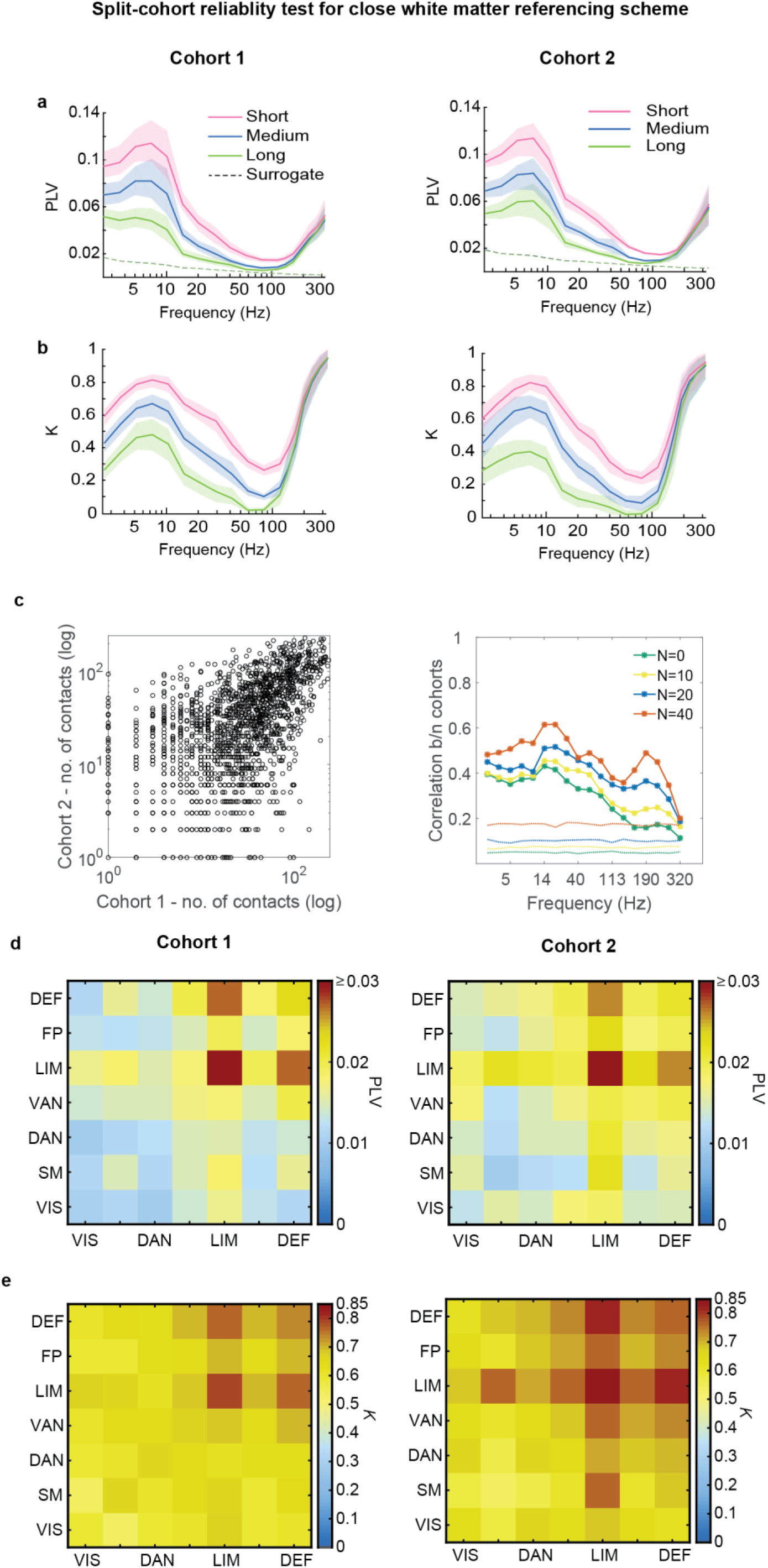
Strength, density and connection patterns of High-gamma phase synchrony are split-cohort reproducible. **a-b,** The mean phase synchrony (PLV) and fraction of significant PLV (*K*) of the two cohorts. **c,** Left: Number of contacts per connection in the Schaefer parcellation for the two cohorts. Right: Spearman correlation between inter-regional functional connection strengths of the two cohorts across all frequencies. Color code: the four solid lines represent the Spearman correlations between connectomes where minimum number of samples to estimate a functional connection was set to 0 (green), 10 (yellow), 20 (blue) and 40 (orange) respectively. Dotted lines: 95^th^ percentile of null distribution of Spearman correlation, where the null distribution was generated by estimating Spearman correlation of randomly resampled (N = 1000, without replacement) versions of the connectomes of the two cohorts. **d-e,** Mean PLV and *K* between Yeo functional systems of the two cohorts.

**Figure S5.**
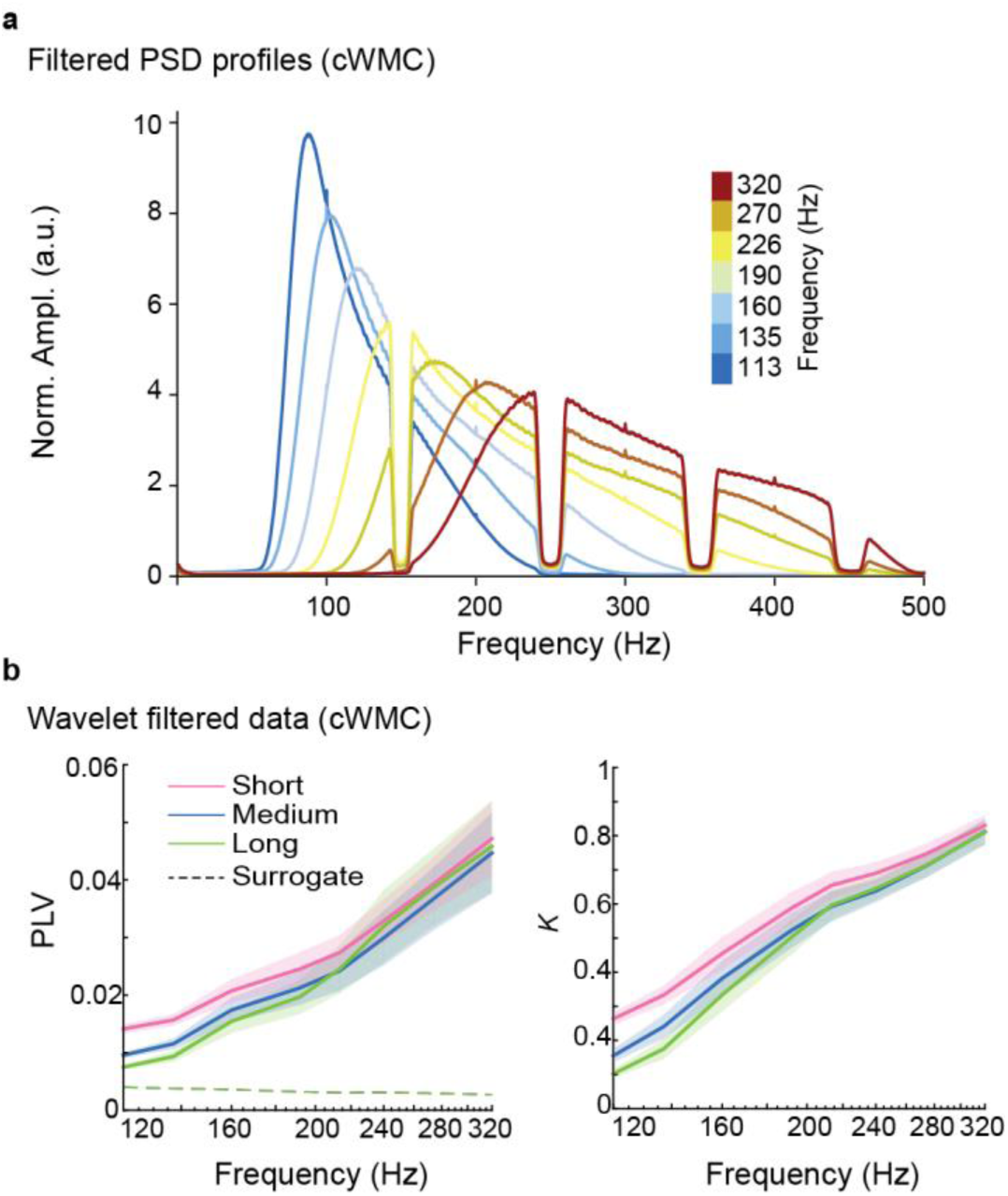
Amplitudes of slower rhythms do not confound HG activity. **a,** The power spectral densities of band-pass filtered data averaged across subjects. Colors represent the central frequency of the band pass. **b,** Mean PLV and connection density *K* for phase synchrony that has been estimated from CW data filtered with Morlet wavelets.

**Figure S6.**
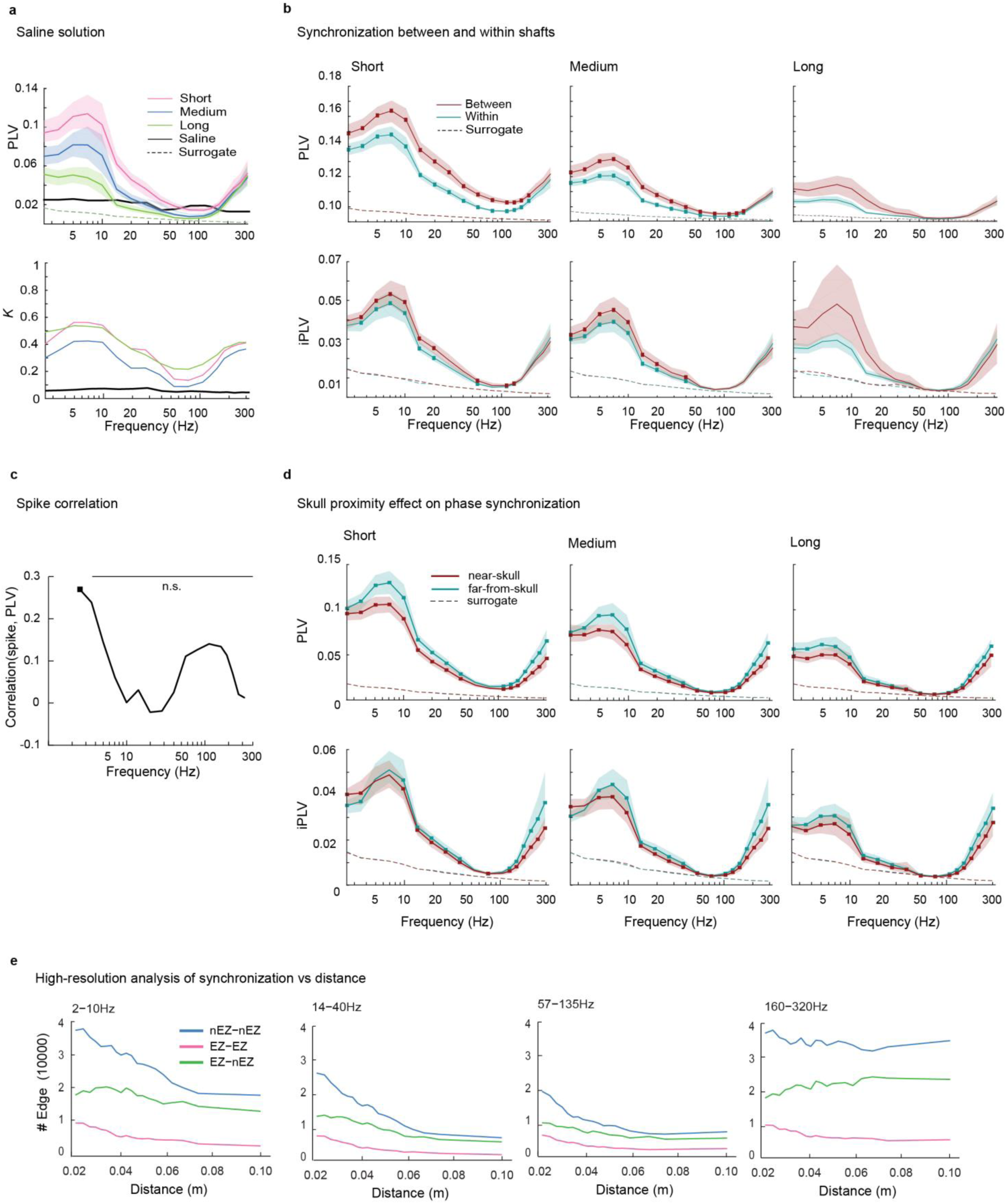
Additional analyses to rule out technical and pathological confounds. **a,** Mean PLV and fraction of significant PLV (*K*) from 10 subjects is compared with that estimated from two electrodes with 18 contacts submerged in a saline solution. **b,** In patients’ brain, mean PLV between contacts from different shafts (red) is larger than for contacts from mean PLV within the same shaft (azure) for all distance ranges. Shaded areas: confidence limits (two-tail; *p* < 0.05) from bootstrapped (N = 100, with replacement) population variance. Square markers indicate significant differences (two-tail permutation test, N = 100, *p* < 0.05). **c,** The Pearson correlation coefficient between edge strengths (PLV) and frequency of inter-ictal events as a function of frequency. Markers: significant correlation (*p* < 0.05, uncorrected). **d,** PLV and iPLV for near-skull (red) and far-from-skull (azure) contacts are reported as a function of frequency for short, medium and long distances. Shaded areas represent confidence limits and square data points represent significant group differences as in **b**. **e,** The mean number of available edges as a function of distance for the computation of distance-specific PLV (see Fig. 7) at different frequency ranges.

**Figure S7.**
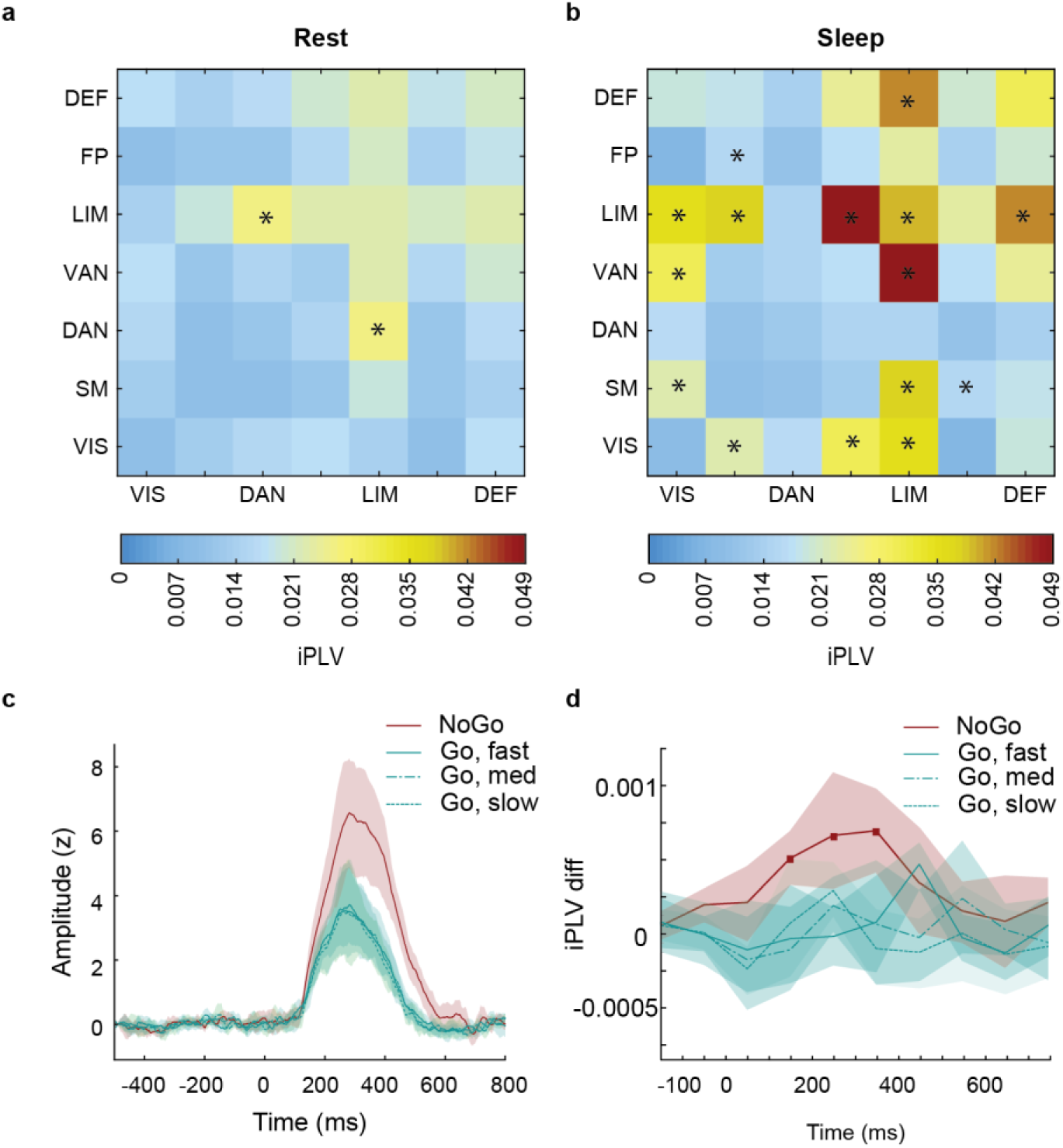
HG Phase synchrony is enhanced during slow wave sleep and visuomotor inhibition. **a-b,** mean iPLV matrices (across HGA frequencies) between Yeo systems are different for rest and sleep (dissimilarity permutation test, *p* < 0.001, N = 1,000, one-tailed) conditions. Higher iPLV values for rest compared to sleep are shown with asterisks on the ‘rest’ image, and vice versa for the ‘sleep’ image (two-tailed t-test, *p* < 0.05, uncorrected). **c,** evoked ERSD for task positive channels in NoGO (red) and Go (azure) conditions. The latter were divided in three groups based on the timing of the behavioral response in fast (plain: 7.3 – 356ms), medium (single dashed: 356 – 459 ms) and slow (double dashed: 459 – 1012 ms) response. Shaded areas represent confidence intervals around mean with 1,000 bootstraps **d,** Evoked iPLV difference from baseline in task positive channels for NoGO (red) and Go (azure) conditions. Shaded areas represent confidence intervals around mean with 1,000 bootstraps. Square markers represent time points where evoked iPLV in NoGo events are significantly different from Go events (permutation test, *p* < 0.05, uncorrected).

**Figure S8.**
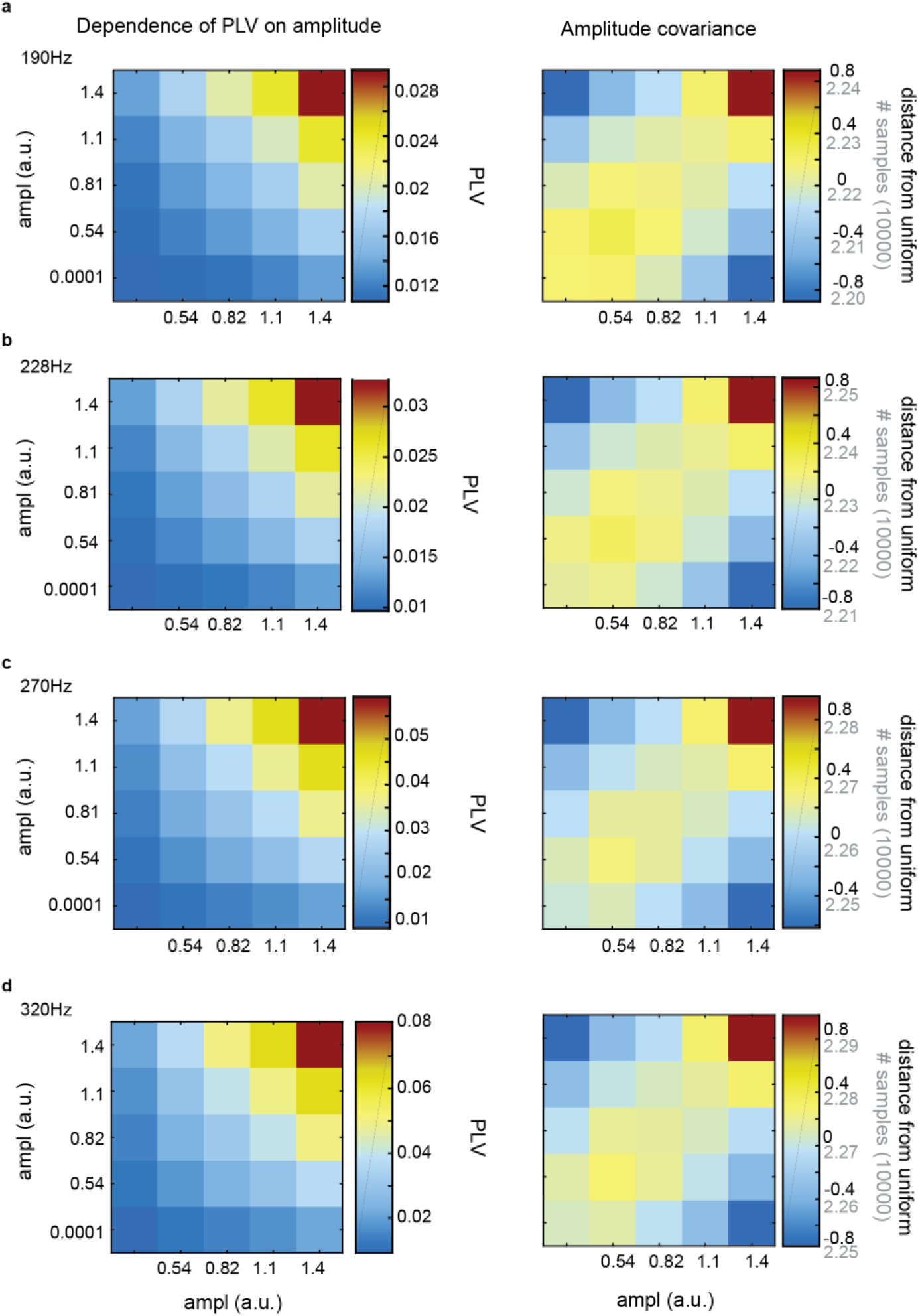
PLV in amplitude bins in High-gamma frequency bands. **a-d,** Joint distribution of moment-to-moment PLV (left column) when two involved channels (x: ch1, y: ch2) demonstrate different amplitude dynamics. Each matrix element is the mean of instantaneous PLV between all significant channel pairs (*p* <0.001 with 100 surrogates) as a function of their moment-to-moment normalized amplitudes. Strength of moment-to-moment amplitude correlation (Right column). Distribution of instantaneous phase samples (light grey) in each amplitude bin and its distance from uniform distribution (black) of samples (*i.e.*, null-hypothesis for the absence of moment-to-moment amplitude correlation). 190Hz, 226Hz, 270Hz, and 320Hz.

**Figure S9.**
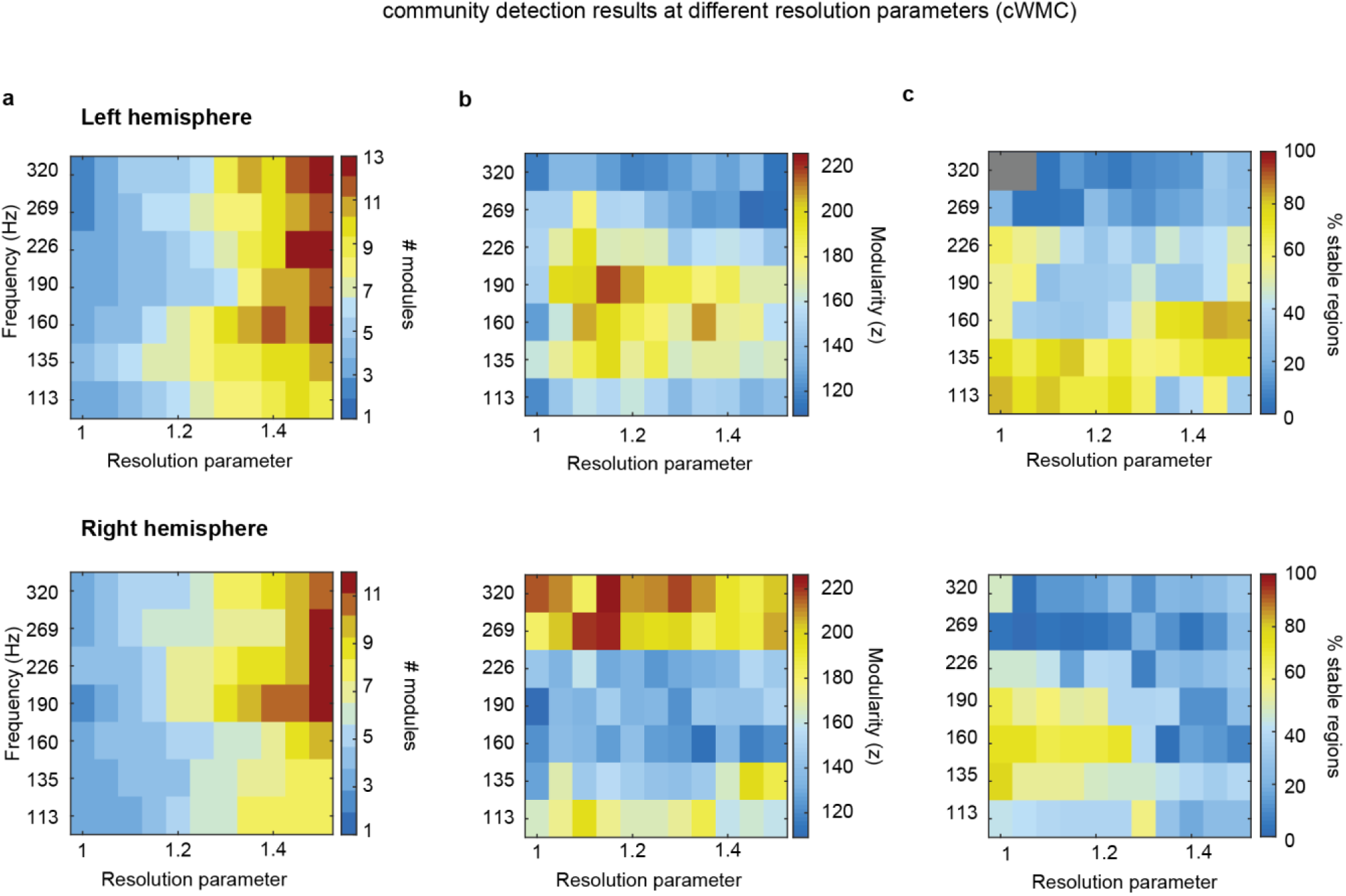
Stable community structure in high-gamma frequency bands across a range of resolutions. **a,** Number of modules for left (top-row) and right (bottom row) hemisphere as a function of resolution parameter γ in high-gamma frequency ranges. **b,** Normalized (z-score) modularity as a function of resolution parameter γ. All frequency and γ combinations yielded significant modularity (permutation test, *p*<0.05, N=100, one-tailed). **c,** Percentage of stable regions for high-gamma bands as a function of resolution parameter γ. Grayed values represent non-significant (bootstrapping test, *p* < 0.05, N=100, one-tailed) combinations.

| ID | Ez Location | age | duration | Sex | Drugs | Outcome |
| --- | --- | --- | --- | --- | --- | --- |
| 1 | Right mesial frontal | 21 | 15:01 | M | CBZ 600 mg- LVT 1500 mg | IB (8 years) |
| 2 | Right temporal insular | 26 | 9:53 | M | CBZ 1200 mg- PRIMIDONE 750 mg- clonazepam 10 mg | IA (37 months) |
| 3 | Right temporal | 38 | 10:29 | M | CBZ 1200 mg- dintoina 450mg lacosamide 300mg clonazepam 10 mg | --- |
| 4 | Left temporal-parietal | 38 | 10:51 | F | FENOBARBITAL 100 MG TOPIRAMATO 100MG LEVETIRACETAM 3000 MG | IA (25 months) |
| 5 | Left temporal-insular | 24 | 10:48 | M | OXC 600 mg- lacosamide 400 mg | IA (15 months) |
| 6 | Right temporo-insular | 40 | 10:33 | F | LBT 1000mg, lacosamide 350mg sertralina 50 mg lorazepam 1mg | IIIA(32 months) |
| 7 | Left temporal- orbital* | 38 | 11:22 | M | CBZ 1200 mg- levetiracetam 3500mg - topamax 200 mg | IA (3 years ) |
| 8 | precuneus | 19 | 11:30 | M | OXC 600 mg- lacosamide 400 mg | --- |
| 9 | functional epilepsy | 20 | 13:13 | F | CBZ 1200 mg - levetiracetam 1500 mg | --- |
| 10 | Right occipito-temporal parietal | 20 | 10:23 | M | lamotrigina 400 mg, levetiracetam 3000mg, lacosamide 400 mg | IA(3 years) |
| 11 | Right temporal-occipital | 41 | 10:03 | M | fenitoina 200mg, lacosamide 300mg | IA (6 months) |
| 12 | Left temporal | 35 | 10:05 | M | topiramato 200mg, CBZ 900mg | IA (12 months) |
| 13 | Rigth anterior temporal | 28 | 10:23 | M | CBZ 1200 mg | IIA( 66 months ) |
| 14 | Temporo-hip* | 36 | 11:26 | F | CBZ 1400 mg levetiracetam 3000 mg | IIA( 6 months) |
| 15 | Left anterior frontal* | 40 | 10:01 | M | levetiracetam 2750mg CBZ 800mg primidone 750mg | IA (12 months) |
| 16 | Left anterior frontal* | 39 | 9:58 | M | oxCarbazepina 1800 mg, clobazam 20 mg, | IA(36 months) |
| 17 | Rigth fronto-temporal-insular | 24 | 10:03 | F | CBZ 1000mg, clobazam 20 mg, lomotrigina 200 mg | IVA(24 months) |
| 18 | Left parieto-opercolo-insular | 31 | 11:09 | F | CBZ 1200 mg, clobazam 40 mg, fenobarbital 75mg | IVA(12 months) |
| 19 | Right temporal perisilvian | 34 | 10:02 | F | Fenobarbital 150 mg, lacosamide 400 mg, clobazam 10mg | IIC(38 months) |
| 20 | Right perisilvian insular | 17 | 10:36 | M | CBZ 800 mg, lamotrigina 400mg | IVA( 26 months) |
| 21 | Right tempo-parieto-occipital | 36 | 10:09 | M | oxCarbazepina 1200mg, fenobarbital 150mg, valproato 1000mg | IA( 35 months) |
| 22 | --- | 32 | 14:40 | F | CBZ 700 mg | --- |
| 23 | Left temporal antero-mesial | 32 | 11:26 | M | CBZ 1200 mg levetiracetam 750 mg | IA (61 months) |
| 24 | Right fronto-centro-insular | 33 | 10:11 | M | CBZ 800mg, lacosamide 800mg, zonisamide 250 mg | IIA (38 months) |
| 25 | Left temporal | 21 | 10:07 | F | levetiracetam 1750 mg, lacosamide 400mg, valproato 1000mg | IA( 24 months) |
| 26 | Right parietal | 23 | 1:54 | M | levetiracetam 3000mg, CBZ 1000mg lacosamide 500mg | IA (24 months) |
| 27 | termo-coagualued multiple sites | 46 | 10:02 | M | CBZ 1200mg, fenobarbital 100mg | IA (24 months) |
| 28 | Right frontal | 20 | 10:48 | F | valproato 800mg, clobazam 10mg | IIA (36 months) |
| 29 | Right mesial frontal | 21 | 16:55 | M | CBZ 800 mg, levetiracetam 3000 mg, nitrazepam 1,5mg | IIIA (13 months) |
| 30 | Rigth fronto central | 22 | 10:10 | M | amotrigina 400 mg, leveracitam 2000mg | IA(24 months) |
| 31 | Right frontal | 20 | 10:33 | M | CBZ 600mg, rufinamide 1500mg, | IVA(13 months) |
| 32 | Right frontal | 44 | 10:05 | F | CBZ 1200mg, zonisamide 400 mg, fenobarbital 1000 mg | IC (24 months) |
| 33 | --- | 17 | 11:13 | M | CBZ 300 mg, | --- |
| 34 | --- | 14 | 10:05 | M | levetiracetam 1500 mg, clobazam 5mg | --- |
| 35 | Right antero mesial temporal | 30 | 10:19 | F | oxcarbazepina 2000mg, fenobarbital 150mg | IIA (36 months) |
| 36 |  | 24 | 10:01 | M | CBZ 16000 mg, levetiracetam 4000 mg | --- |
| **37** |  | 29 | 9:53 | F | levetiracetam 3000 mg | --- |
| **38** | Right orbito temporal | 29 | 9:56 | F | zonisamide 400 mg, levetiracetam 750 mg, CBZ 1400 mg | IA (62 months) |
| **39** |  | 45 | 10:10 | F | lacosamide 500 mg, valproato 1000 mg, zonisamide 200 mg | --- |
| **40** | termo-coagualed multiple sites | 34 | 10:00 | F | CBZ 1000mg ,levetiracetam 2500mg | IA( 12 months) |
| **41** | termo-coagualed multiple sites | 50 | 10:00 | M | levetiracetam 2000mg, lacosamide 600mg | IIA( 6months) |
| **42** | Left occipital | 17 | 10:03 | F | CBZ 1200 mg, levetiracetam 1500 mg, lacosamide 300 mg | IB (49 MES) |
| **43** | Right temporal | 44 | 10:03 | F | topiramato 300 mg, oxcarbamazepina 1200 mg | IIA (50 months) |
| **44** |  | 27 | 9:58 | M | CBZ 800 mg, lamotrigina 200 mg | --- |
| **45** |  | 46 | 10:01 | M | CBZ 1200 mg, levetiracetam 3000 mg, lacosamide 150mg, clobazam 20mg, | --- |
| **46** | Left cingulate | 30 | 10:02 | M | oxcarbamazepina 1800 mg, topiramato 200 mg, levetiracetam 3000 mg, clobazam 10mg | IA (16 months) |
| **47** | Right anterior frontal | 28 | 11:07 | M | CBZ 1000 mg, levetiracetam 1000 mg | IIIA (61 months) |
| **48** | Right temporo-parieto-perisylvian* | 27 | 10:01 | F | topiramato 200 mg, lamotrigina 200 mg | IA (5 years) |
| **49** | Right antero mesial temporal | 42 | 11:28 | F | lacosamide 500 mg, | IB (36months) |
| **50** | Left parieto-temporal | 15 | 10:08 | M | CBZ 900mg | IA (6 months) |
| **51** | Right temporo-opercular* | 37 | 10:02 | M | CBZ 900mg, levetiracetam 3000mg | IVA (12 months) |
| **52** | Left frontal | 30 | 10:16 | F | CBZ 1200mg, lamotrigina 200 mg, cllobazam 20mg | IA(5 years) |
| **53** | Left frontal | 15 | 10:00 | F | levetiracetam 1250 mg, oxcarbamazepina 1200 mg | IA (4 years) |
| **54** | termo-coagualed multiple sites | 41 | 10:04 | M | levetiracetam 3000 mg, lacosamide 400 mg | IA (2 years) |
| **55** | Right temporo-occipital | 37 | 10:07 | M | lamotrigina 600mg, levetiracetam 2000 mg | IA (2 years) |
| **56** | Right temporal | 29 | 10:20 | M | CBZ 1400 mg, levetiracetam 3000 mg, clobazam 10 mg | IA (31 months) |
| **57** | Left operculo insula | 10 | 10:02 | F | CBZ 800 mg | IIIA (34 months) |
| **58** | Right temporo frontal | 40 | 10:02 | F | lamotrigina 600mg, clobazam 20mg, fenitoina 500 mg | IA (13 months) |
| **59** | Left temporo insula opercular | 29 | 11:04 | F | CBZ 1800 mg, clobazam 20 mg | IA (24 months) |
| **60** | Right temporal | 27 | 11:15 | M | lamotrigina 400 mg, topiramato 400 mg | IA (24 months) |
| **61** |  | 26 | 9:54 | M | fenitoina 400mg, topiramato 500 mg | --- |
| **62** | Left temporal | 17 | 10:15 | M | oxcarbamazepina 1500 mg, clobazam 20 mg, levetiracetam 2500 mg | IA (36 months) |
| **63** | Right mesial temporal | 25 | 11:03 | F | topiramato 75mg, CBZ 1500 mg | IA (12 months) |
| **64** | Nodular heterotopy* | 24 | 10:11 | F | CBZ 1000 mg, levetiracetam 500 mg, clobazam 20 mg | IVA (12 months) |
| **65** | Left temporo perisylvian | 37 | 9:54 | F | CBZ 1200mg, lamotrigina 550mg, | IIIA( 12 months) |
| **66** | Left antero-mesial temporal | 32 | 10:30 | F | clobazam 20mg, fenobarbital 45mg | IIA (55 months) |
| **67** | Right temporo occipital | 44 | 10:02 | M | CBZ 800mg, levetiracetam 3000mg, fenobarbital 125mg | IA (24months) |

